# Growth cone macropinocytosis of neurotrophin receptor and neuritogenesis are regulated by Neuron Navigator 1

**DOI:** 10.1101/2022.01.31.478516

**Authors:** Regina M. Powers, Ray Daza, Alanna E. Koehler, Julien, Barbara Calabrese, Robert F. Hevner, Shelley Halpain

## Abstract

Neuron Navigator 1 (Nav1) is a cytoskeleton-associated protein expressed during brain development that is necessary for proper neuritogenesis, but the underlying mechanisms are poorly understood. Here we show that Nav1 is present in elongating axon tracts during mouse brain embryogenesis. We found that depletion of Nav1 in cultured neurons disrupts growth cone morphology and neurotrophin-stimulated neuritogenesis. In addition to regulating both F-actin and microtubule properties, Nav1 promotes actin-rich membrane ruffles in the growth cone, and promotes macropinocytosis at those membrane ruffles, including internalization of the TrkB receptor for the neurotrophin BDNF (brain-derived neurotropic factor). Growth cone macropinocytosis is important for downstream signaling, neurite targeting, and membrane recycling, implicating Nav1 in one or more of these processes. Depletion of Nav1 also induces transient membrane blebbing via disruption of signaling in the Rho GTPase signaling pathway, supporting the novel role of Nav1 in dynamic actin-based membrane regulation at the cell periphery. These data demonstrate that Nav1 works at the interface of microtubules, actin, and plasma membrane to organize the cell periphery and promote uptake of growth and guidance cues to facilitate neural morphogenesis during development.

## INTRODUCTION

Mammalian brain development requires a series of steps coordinated by intra- and extra-cellular cues. Integration of these cues by developing neurons results in morphological changes guiding the maturation of newborn neurons and their ultimate integration into neuronal networks. Actin and microtubules (MTs) of the neuronal cytoskeleton, regulated by cytoskeleton-associated proteins, undergo major rearrangements to drive neuronal morphogenesis throughout development.

One group of critical cytoskeleton-associated proteins are microtubule plus-end tracking proteins (+TIP proteins). +TIPs localize to the dynamic, growing end of MTs and form complexes there, many anchored by end-binding proteins (EBs), which bind directly to the microtubule plus end (Akhmanova and Steinmetz, 2008). In addition, actin filaments and MTs can interact to deliver molecules to and from the cell periphery and initiate membrane remodeling in response to signals, and +TIPs have been shown to mediate certain types of MT-F-actin interactions (Geraldo *et al*., 2008; Preciado Lopez *et al*., 2014; Henty-Ridilla *et al*., 2016). Such precisely regulated cytoskeleton dynamics are especially important in the neuronal growth cone, a specialized cytoskeleton-rich structure that responds to extracellular cues and directs neurite outgrowth (Dehmelt and Halpain, 2004; Geraldo *et al*., 2008; Dent *et al*., 2011; Preciado Lopez *et al*., 2014; Pacheco and Gallo, 2016).

The growth cone comprises cytoskeletal structures containing both MTs and actin, including microtubule arrays, filopodia, actin arcs, and F-actin membrane ruffles (Suter and Forscher, 2000; Dent *et al*., 2011), and their regulated dynamics are essential for growth cone responses to external cues and proper neurite outgrowth and guidance (Lin and Forscher, 1995; Mitchison and Cramer, 1996; Lowery and Van Vactor, 2009; Dent *et al*., 2011; McCormick and Gupton, 2020). The role of growth cone F-actin membrane ruffles in growth cone behavior, however, remains largely unexplored. In non-neuronal cells, membrane ruffles influence cell migration, morphogenesis, and endocytosis (Radhakrishna *et al*., 1999; Borm *et al*., 2005; Lim and Gleeson, 2011). Notably, Bonanomi *et al* found that growth cone membrane ruffles are sites of bulk-endocytosis, and that this endocytosis is important for neuritogenesis (Bonanomi *et al*., 2008). However, the mechanisms that regulate growth cone membrane ruffles and their functional impact are largely unexplored.

The Neuron Navigators (Nav1, Nav2, and Nav3) are a group of +TIP proteins functioning in development and tissue morphogenesis. All three proteins are large (>200kDa), and contain multiple domains including a microtubule binding domain and a AAA ATPase domain whose function is poorly understood (Maes *et al*., 2002). Nav1 is enriched in the developing nervous system (Martinez-Lopez *et al*., 2005), and concentrates at neurite tips (van Haren *et al*., 2014). Both Nav1 and Nav2 have been implicated in neuritogenesis (Martinez-Lopez *et al*., 2005; Muley *et al*., 2008; van Haren *et al*., 2009; Marzinke *et al*., 2013; van Haren *et al*., 2014). Nav1 regulates neuritogenesis in neuroblastoma cells via binding to the Rho GEF Trio (van Haren *et al*., 2014). Nav1 might also facilitate MT-actin crosstalk via direct F-actin binding (Sanchez-Huertas *et al*., 2020). While these studies provide insight into the role of Nav1 in neural development, many of the underlying mechanisms remain to be elucidated. In this paper, we investigate the critical role of Nav1 in neuritogenesis in multiple model systems, describe its selective enrichment in and function in growth cone membrane ruffles, and identify two novel roles of Nav1 in regulating the cytoskeleton. These novel roles include the regulation of cortical actin-plasma membrane connections via the Rho/Rac pathway, and a key role in macropinocytic uptake of neurotrophin receptors in the nascent growth cone via regulation of membrane ruffles.

## RESULTS

### Nav1 is enriched in pathfinding axons *in vivo*

Nav1 mRNA is enriched in the developing CNS (Martinez-Lopez *et al*., 2005), but thus far the *in vivo* localization of Nav1 protein regionally within the brain has not been described. We characterized Nav1 immunoreactivity in embryonic mice at E12.5 and E14.5. As expected, Nav1 mRNA was enriched throughout the developing brain and enriched in the majority of neuronal cell bodies of differentiated neurons (Visel *et al*., 2004, Figure 1A-B). Within the developing neocortex, immunoreactivity for Nav1 was low in the progenitor-containing ventricular zone (VZ) and subventricular zone (SVZ) (Figure 1C). However, Nav1 immunoreactivity was abundant in the intermediate zone (IZ) and cortical plate (CP) (Figure 1C). In addition to enrichment in cell bodies (Figure 1D,E), Nav1 immunoreactivity was enriched in several developing axon tracts, including the lateral olfactory tract, the stria medullaris thalami, and the internal capsule (Figure 1C). Within these axon tracts and within the IZ, Nav1 colocalized with multiple axonal markers, including antibody SMI-31, which labels neurofilaments (Figure 1F- K), antibody to growth associated protein-43 (GAP-43) (Figure 1L-N), and antibody TuJ1, which labels neuron-selective betaIII-tubulin (Binder *et al*., 1985) (Figure 1O,P). Enrichment of Nav1 in these axon tracts is consistent with a role in axon outgrowth or pathfinding *in vivo*.

**Figure 1.**
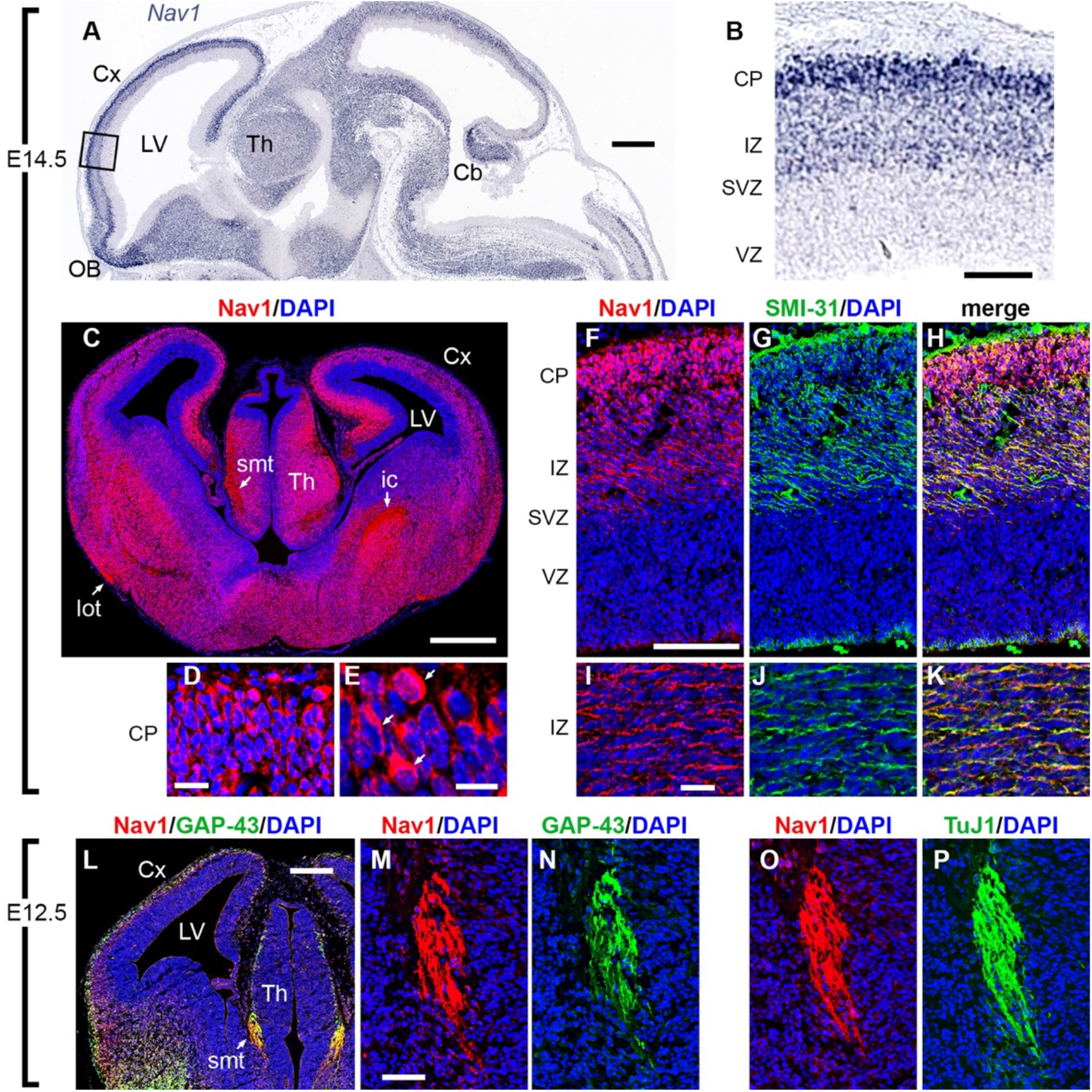
Nav1 is localized in neuronal cell bodies and growing axons in developing mouse brain. (A, B) Nav1 mRNA, detected by in situ hybridization (ISH) on E14.5 (Genepaint, set ID EH1164), was widely expressed in differentiating neurons, but not in progenitor cells around the ventricles. In cerebral cortex (boxed area in A, shown at higher magnification in B), Nav1 mRNA was not detected in the progenitor-containing ventricular zone (VZ) and subventricular zone (SVZ), but was highly expressed in the intermediate zone (IZ) and cortical plate (CP), where postmitotic neurons are located (Genepaint, set ID EH1164). (C-E) Nav1 protein on E14.5 was expressed in neuronal cell bodies, but was also highly enriched in axon tracts including the stria medullaris thalami (smt), lateral olfactory tract (lot), and internal capsule (ic). Higher magnification views of the CP in D and E demonstrate Nav1 protein in many cell bodies (arrows in E). (F-K) Double immunofluorescence for Nav1 and neurofilament heavy chain (antibody SMI-31), a marker of axons, confirmed the presence of Nav1 in axons coursing through the IZ. (L-N) On E12.5, double immunofluorescence for Nav1 and GAP-43, a marker of growth cones, demonstrated extensive colocalization, notably in growing axons of the smt, shown at higher magnification in M-N. (O, P) Double immunofluorescence for Nav1 and class III b- tubulin (antibody TuJ1), a marker of neurons and their processes, confirmed the presence of Nav1 in smt axons. Plane of section: sagittal for A, B; coronal for C-P. Scale bars: A, 500 μm; B, 100 μm; C, 500 μm; D, 20 μm; E, 10 μm; F, 100 μm (for F-H); I, 20 μm (for I-K); L, 200 μm; M, 50 μm (for M-P). Cx = Cortex, LV = Lateral Ventricle, Th = Thalamus, OB = Olfactory Bulb, Cb = Cerebellum.

### Nav1 accumulates in subcellular regions of morphological change

Together with previous studies (Martinez-Lopez *et al*., 2005; van Haren *et al*., 2009; van Haren *et al*., 2014; Sanchez-Huertas *et al*., 2020), the observation of Nav1 expression in axonal tracts during the phase of outgrowth and cue-driven guidance *in vivo* suggests that Nav1 is involved in neurite development. We therefore assessed the localization of endogenous Nav1 in multiple neural cell types. We used dissociated hippocampal neurons cultured from embryonic rat brain, SH-SY5Y human neuroblastoma cells, and human iPSC-derived neurons as model systems to further investigate the functional role of Nav1 in developing neurons. These cell models recapitulate in culture many of the key morphological changes that neurons undergo during neuritogenesis *in vivo*, and are amenable to detailed sub-cellular studies and time-lapse observation (Merrill *et al*., 2002; Muley *et al*., 2008; Agholme *et al*., 2010).

In newly-plated primary hippocampal neurons that had yet to extend processes, endogenous Nav1 was distributed in a punctate pattern throughout the cell. However, clusters of Nav1 puncta were especially enriched at peripheral locations lying within several micrometers of the plasma membrane (Figure 2A, Stage 1.1). Clusters of Nav1 puncta were especially enriched at peripheral locations, which are often sites of symmetry breakage where *in vitro* growth cone formation and neurite initiation commonly occur in this model (Dehmelt and Halpain, 2004; Dehmelt *et al*., 2006). Neurons transfected with GFP-Nav1 displayed an even greater clustering of GFP-Nav1 in growth cones and their precursors, and time-lapse imaging confirmed that GFP- Nav1 was concentrated within subcellular domains that are presumptive sites of neurite initiation (Supp. Figure 1). Regions enriched with Nav1 redistributed as the cell margin dynamically segmented and coalesced into nascent growth cones (Figure 2A, Stage 1.2-1.3). Nav1 enrichment was evident in all identified growth cones of minor neurites (Figure 2A, Stage2). Growth cone enrichment of Nav1 continued throughout the early stages of neurite elongation, and persisted as one of the minor neurites began differentiating into an axon (Figure 2A, Stage 3). This pattern of Nav1 localization is consistent with a potential role in directing multiple stages of neuritogenesis, including neurite initiation, elongation, and axon guidance. Experiments demonstrated that clusters of Nav1 were present in growth cone regions containing enriched immunoreactivity to the actin binding protein drebrin (Figure 2B), which we used as an F-actin marker compatible with the methanol fixation method that optimizes Nav1 immunoreactivity. In addition, many Nav1 puncta colocalized with tyrosinated tubulin, which labels newly polymerized microtubules (Janke and Bulinski, 2011) (Figure 2C).

**Figure 2.**
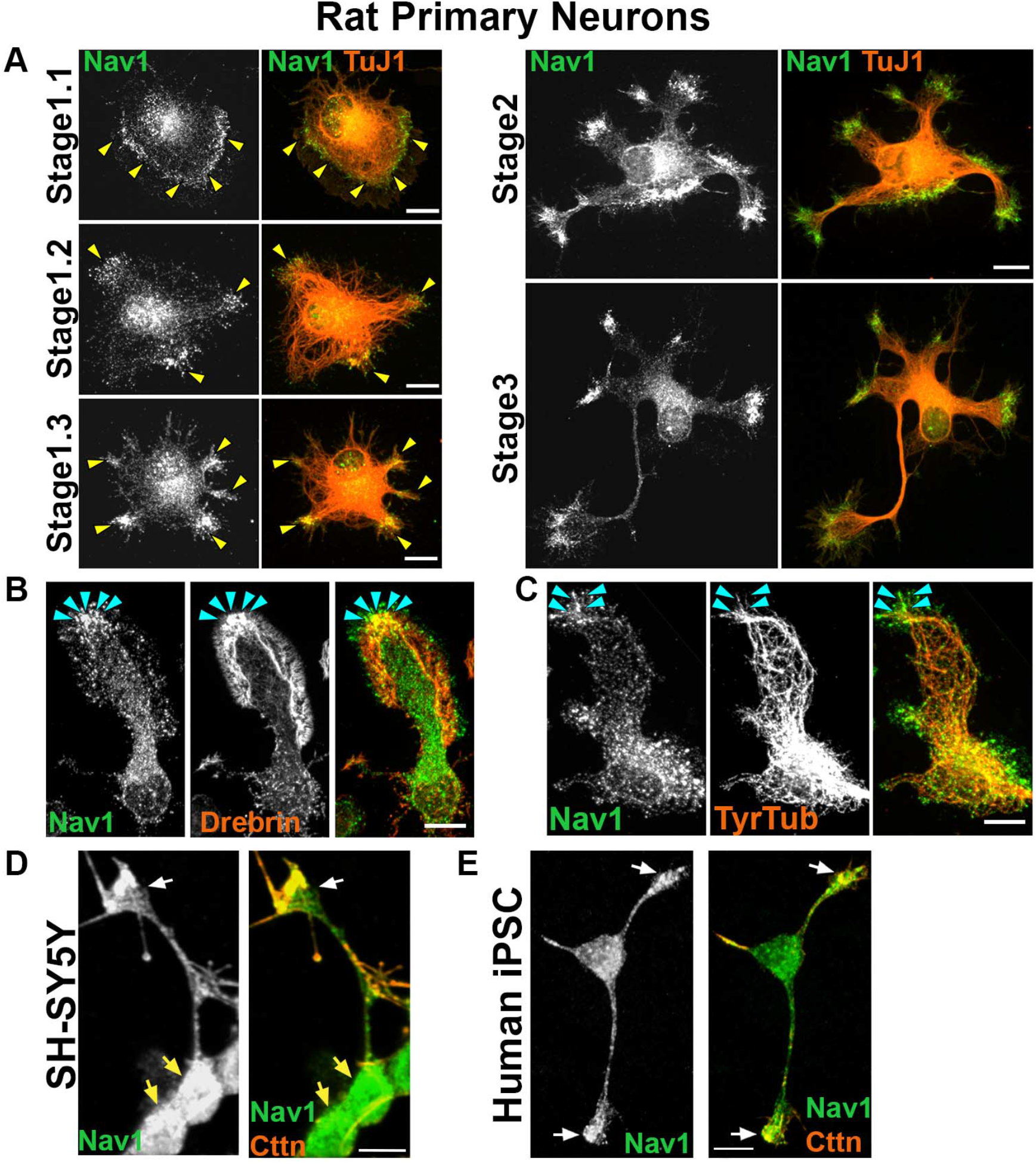
Nav1 is expressed in areas of morphological change. (A) Primary cultures from rat hippocampus were fixed after 3 DIV and immunostained for Nav1 and neuron-specific ß-III tubulin (antibody TuJ1). Representative images of neurons at different stages of neuritogenesis are shown. In Stage 1.1 neurons have a lamellipodia surrounding the cell body and Nav1 is enriched close to the membrane in this area. As the lamellipodium undergoes segmentation (Stage 1.2), Nav1 remains enriched in this location. The segments coalesce into nascent growth cones as the microtubules adopt a parallel organization behind them (Stage 1.3). Nav1 remains similarly clustered in nascent growth cones as minor neurites become established in Stage 2, and one neurite is specified as the axon in Stage 3. (B) Representative image of a growth cone immunostained for Nav1 and the actin binding protein drebrin, highlighting the transition zone of the growth cone. (C) Representative image of growth cone immunostained for Nav1 and tyrosinated tubulin shows that newly-polymerized microtubules are present together with Nav1. (D) Representative image of 3 day BDNF-differentiated SH-SY5Y cells showing enriched Nav1 puncta distal ends of neurite (white arrow indicate growth cone, yellow arrows indicate cell bodies). (E) Representative image of human iPSC-derived neuron showing enriched Nav1 in distal ends of neurites (white arrows indicate growth cones). All scale bars = 10μm.

Human SH-SY5Y cells stop dividing, differentiate into neuronal-like cells, and emit neurites after sequential stimulation with retinoic acid and brain-derived neurotrophic factor (BDNF) (Encinas *et al*., 2000). Similar to the rat hippocampal neurons, differentiated SH-SY5Y cells displayed enriched clusters of Nav1 puncta in the distal ends of neurites (Figure 2D), although the relative degree of enrichment was lower than what was observed in hippocampal neurons. Finally, we also observed enriched Nav1 in growth cones of human iPSC-derived neurons (Figure 2E). Taken together, these data show that Nav1 is present in developing human and rodent neurons and accumulates in areas of morphological change as the nascent neurites are forming.

### Nav1 deficiency disrupts neuritogenesis

Previous studies have shown that knockdown of Nav1 in a mouse neuroblastoma cell model disrupts neuritogenesis (van Haren *et al*., 2014). To address whether Nav1 plays a similar role in primary neurons, we performed knockdown experiments in freshly dissociated hippocampal neurons (Figure 3A). Cells were electroporated with control vector or Nav1-shRNA just prior to plating, and examined at 3 days in vitro (3DIV). Nav1 shRNA induced a ∼50% reduction in neuronal Nav1 expression compared to control shRNA (Supp. Fig 2A). First, we analyzed the effect of Nav1 silencing on early morphogenesis by classifying neurons into three stages (Dotti *et al*., 1988): stage 1 cells have no identifiable neurites; stage 2 cells have one or more minor neurites but lack a defined axon; stage 3 cells have one or more axons. The proportion of cells in stage 1 was significantly higher in Nav1-depleted neurons, with no significant effect on the proportions of cells in stages 2 and 3 (Figure 3B). This indicates that Nav1 directs neuronal morphology from the earliest morphological steps.

**Figure 3.**
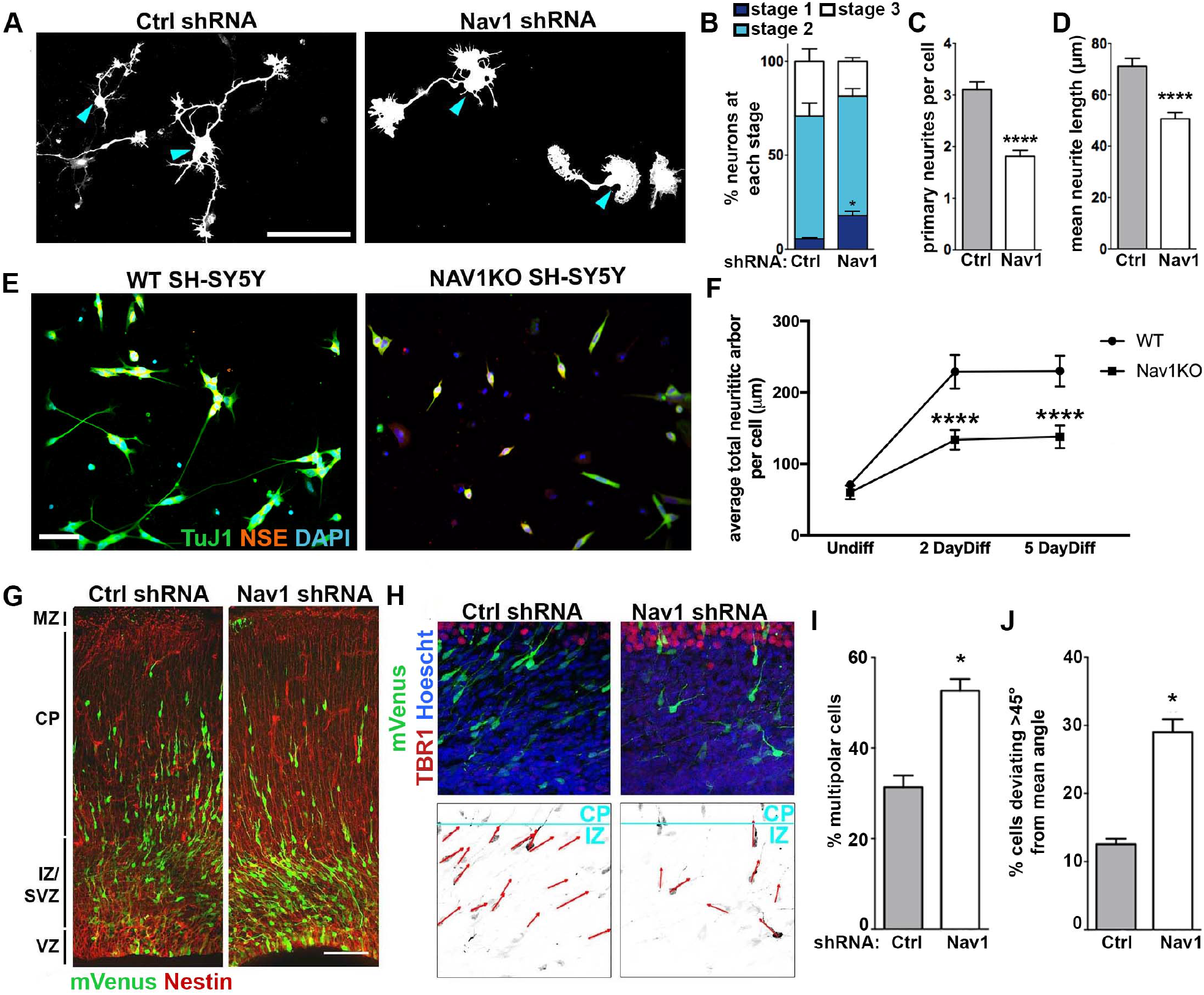
Nav1 deficiency disrupts neuritogenesis. (A) Hippocampal neurons were electroporated on the day of plating to express GFP as transfection marker and either Control shRNA (Ctrl shRNA) or Nav1 shRNA. Neurons were fixed after 3 DIV and stained for GFP and ß-III-tubulin to specifically identify neurons. Representative GFP images are shown; blue arrowheads indicate the cell body. Scale bar = 100μm (B) Quantification of neurite growth in Ctrl and Nav1 shRNA electroporated dissociated hippocampal neurons: Neurons were classified into different stages as described in Methods. Stacked histogram showing that knock-down of Nav1 increases the percentage of cells in stage 1. Statistical analysis: Two-way ANOVA with Bonferroni post hoc comparisons, *p<0.05.; n = 3 experiments; Control shRNA = 240 cells, Nav1 shRNA = 352 cells (C) Nav1- supressed neurons show a decrease in the number of primary neurites. Statistical analysis: Mann-Whitney test ****p<0.0001. Same n as (B). (D) Nav1-suppressed neurons show decreased mean neurite length. Mann-Whitney test ****p<0.0001. Same n as (B). (E) Representative images showing WT and Nav1KO SH-SY5Y cells after 2 days of BDNF differentiation as described in Methods. Scale bar = 50μm (F) Nav1KO cells have reduced neurite growth over time. Neurite length quantified as described in methods. Statistical analysis: Kruskal-Wallis, Dunn’s multiple comparisons ****p<0.0001; n = 3 experiments; Undiff WT = 226 images, Undiff Nav1KO = 238 images, 2 day diff WT = 234 images, 2 day diff Nav1KO = 193 images, 5 day diff WT = 232 images, 5 day diff Nav1KO = 175 images; (G) Typical example of neuronal morphology and orientation of cells electroporated on E15.5 with mVenus (detected using antibody to GFP; green) and either Ctrl shRNA or Nav1 shRNA. Tissue was fixed and counterstained with anti-nestin antibody (red) on E18.5. Scale bar = 100μm. (H) Representative images of Control shRNA and Nav1 shRNA-expressing neurons (green) illustrating the vector (red) used to measure the angle to the pia surface (blue). Tbr1 antibody was used to label the cortical plate. Scale bar = 10μm. (I) Quantification of the effect of Nav1 shRNA on the percentage of cells in the multipolar stage within the IZ/SVZ. Statistical analysis: T test, two tailed, *p<0.05; Control shRNA vector, n = 3 experiments with 6 embryos; Nav1 shRNA vector, n = 3 experiments with 7 embryos. (J) Quantification of the effect of Nav1 shRNA on neuron orientation in the lower IZ and SVZ. Angle of orientation in control vs. shRNA expressing neurons was quantified as described in Methods. Statistical analysis same as (I). All data are expressed as mean +/- SEM.

Consistent with a role in neurite initiation we observed that the average number of primary neurites per cell was significantly reduced by the partial Nav1 knockdown (Figure 3C). The total length of neurite arbor was also modestly but significantly reduced, an effect that could result from reduced neurite initiation, reduced elongation, or both. Because the average neurite length was reduced in the knockdown condition (Figure 3D), and the length of the longest process was also reduced, it seems likely that Nav1 affects neurite elongation as well as neurite initiation.

Many primary hippocampal neurons begin emitting neurites within 4-6 hours of plating, making it difficult to ensure that genetic silencing is uniformly effective prior to the initiation of neuronal morphogenesis. Therefore, we created a Nav1knockout (Nav1KO) line using CRISPR/Cas9 in SH-SY5Y cells to examine the effect of complete Nav1 silencing on differentiation and neuritogenesis (Supp. Figure 2B,C). As described above, differentiated SH-SY5Y cells emit neurites that grow longer over time. We differentiated WT and Nav1KO SH-SY5Y cells, and fixed and assessed them at various times post-differentiation. Nav1KO cells stopped dividing and displayed small neurites positive for TuJ1. However, they showed a significantly lower total neuritic arbor length at both differentiated time points measured, and no difference when undifferentiated (Representative image Figure 3E, quantified in Figure 3F).

### Nav1 deficiency disrupts neuronal morphogenesis *in vivo*

To corroborate the above observations that Nav1 depletion disrupts neuronal morphogenesis, we utilized *in utero* electroporation to introduce Nav1 shRNA and control shRNA constructs co-transfected with mVenus into neural progenitor cells in mouse neocortex at embryonic day E15.5. Labeled cells in the developing cortex were examined 3 days later, at E18.5. In brains of embryos treated with control shRNA, approximately two-thirds of the labeled cells had a bipolar morphology, and the remainder had a multipolar morphology. In contrast, the brains of knockdown embryos showed a significant increase in the fraction of multipolar cells, and a corresponding decrease in bipolar cells (Figure 3G,I). When Nav1 knockdown neurons did display a bipolar morphology, they often appeared to be misoriented. In controls, labeled bipolar neurons within the SVZ and lower IZ had a prominent leading process that was oriented toward the pial surface, roughly perpendicular to the plane of the pial surface toward which it is expected to migrate. Instead, the leading process of Nav1-silenced neurons deviated substantially from the perpendicular and were not consistently oriented in similar directions (Figure 3H,J).

These data suggest that Nav1 is important for the proper orientation of neuronal morphology during cortical cell migration, possibly reflecting a failure to respond to external directional cues. Taken together, the disruption of neural morphogenesis with Nav1 depletion *in vitro* and *in vivo* demonstrate the importance of Nav1 in regulating early neuronal morphogenesis, but the question remained of the role of Nav1 at the growth cone and its influence during development.

### Nav1 and EBs have distinct distributions within the growth cone

We more closely examined the distribution of Nav1 in the growth cone to gain insight into the role of Nav1 there. Where possible we took advantage of the occasional examples of especially large, well-spread growth cones in rat hippocampal cultures, where we could more readily distinguish among the three subcellular domains that have been defined for growth cones: the MT-rich central domain (C-domain) the F-actin-rich peripheral domain (P-domain), and the region between them, the transition zone (T-zone) (Forscher and Smith, 1988). Previous research shows that Nav1 is a +TIP protein that is frequently found on growing microtubule ends containing EB1 and/or EB3, and that Nav1 and EB binding is mediated by the Nav1 microtubule binding domain (van Haren *et al*., 2009; Sanchez-Huertas *et al*., 2020). Within neuronal growth cones, advancing microtubules often extend from the C-domain across the T-zone, and frequently invade filopodia in the P-domain (Schaefer *et al*., 2002; Geraldo *et al*., 2008; Pacheco and Gallo, 2016). We observed that Nav1 immunoreactivity was often (but not always) associated with the tip of filopodial microtubules, as expected for a +TIP protein (Figure 4A; yellow arrowheads). Such microtubules were recently polymerized and presumably dynamic, since they were immunopositive for tyrosinated tubulin (Figure 4B). However, Nav1 puncta could also be observed along the length of some microtubules (Figure 4A; cyan arrowheads), as well as in filopodia that were devoid of detectable microtubules (Figure 4A; white arrowheads). Thus, it appears that Nav1 is neither an obligatory nor an exclusive member of +TIP complexes in the growth cone.

**Figure 4.**
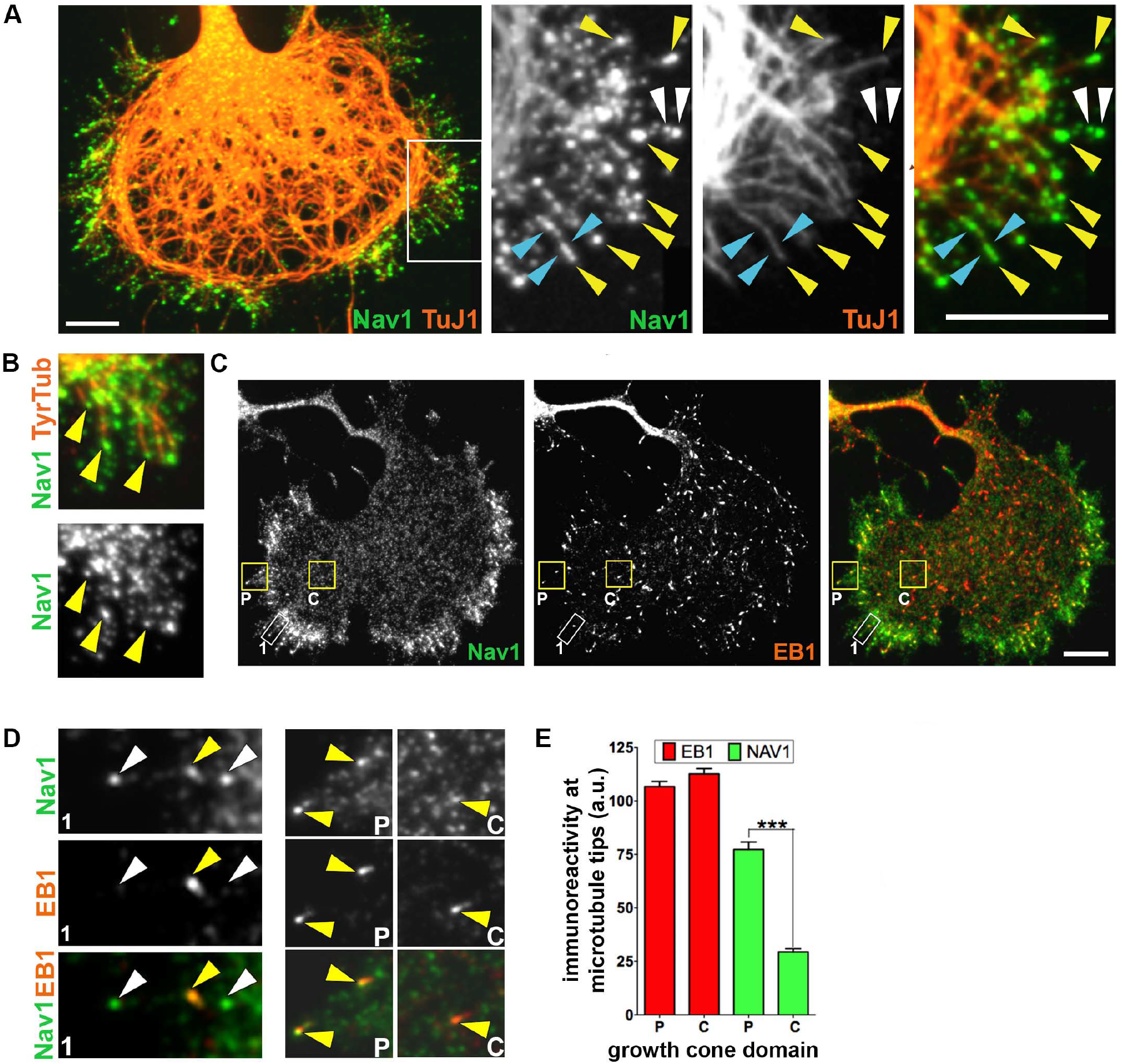
Nav1 is a +TIP protein that associates with EB1 in the growth cone. (A) Growth cone stained for Nav1 and neuron-specific marker TuJ1 (ß -III tubulin). The white box indicates the region enlarged in three panels to the right. Note that Nav1 puncta are present at the tips of microtubules in the P-domain and T-zone. Yellow arrowheads indicate NAV1 puncta at the tips of individual microtubules; blue arrowheads indicate Nav1 puncta arrayed along microtubules; white arrowheads indicate Nav1 puncta not colocalized with detectable microtubules. Scale bar = 10μm (B) Nav1 localizes to the tips of newly polymerized microtubules (arrowheads) labeled with antibody to tyrosinated tubulin. (C) Growth cone immunostained for Nav1 and EB1. Note that the overall distribution of Nav1 is different from that of EB1; NAV1 puncta accumulate within the T-zone, while EB1 puncta distribute more uniformly. Boxed regions are shown at higher magnification in (D). Scale bar = 10μm (D) Box 1 is zoomed region outlined by white rectangle, Box P (P-domain) and Box C (C-domain) outlined in yellow rectangles in (C). Note that of the three Nav1 puncta in box 1 (arrowheads) only one detectably colocalizes with EB1 (yellow arrowhead). Arrowheads in Box P indicate two intense Nav1 puncta that colocalize with intense EB1 puncta. Single arrowhead in Box C indicates a faint Nav1 punctum that colocalizes with an intense EB1 punctum. (Note that the display settings in zoomed regions are enhanced from those in b.) (E) Quantification of the relative concentration of endogenous EB1 and Nav1 at microtubule tips in the P-domain versus the C-domain. This assay indicates that Nav1 concentration at plus ends is greater in the P-domain and T-zone compared to the C domain. Statistical analysis: Mann-Whitney test; ****p<0.0001; n = 3 experiments; P-domain = 234 comets; C- domain = 243 comets. a.u. = arbitrary units. All data are expressed as mean +/- SEM.

Indeed, endogenous Nav1 showed a distribution pattern in growth cones that was overall very different from that of EB1 and EB3. EB1 immunoreactivity was present as small, often comet-shaped puncta that were uniformly distributed throughout all domains of the growth cone (Figure 4C). In contrast, endogenous Nav1 puncta were relatively enriched within the T-zone (Figure 4C). Similarly, the overall distribution of puncta of ectopic EB3-mCherry was fairly uniform throughout the growth cone, and distinct from that of ectopic GFP-Nav1 (Supp. Fig 3A). In time-lapse imaging EB3-mCherry showed the dynamic, comet-like behavior that is characteristic of +TIP proteins (Perez *et al*., 1999). Individual puncta that contained both EB3-mCherry and GFP- Nav1 moved in concert, and GFP-Nav1 puncta often invaded growth cone filopodia, as described previously for the EBs (Supp. Fig3B). Quantitative image analysis in fixed specimens revealed that endogenous Nav1 displayed a significantly higher coefficient of variation (CV) for its distribution profile across the growth cone compared to that for endogenous EB1 in the same cells (CV for Nav1 = 1.17 ± 0.02; CV for EB1 = 0.57 ± 0.02, n = 91 cells; Mann-Whitney test, p<0.0001), thereby confirming the distinct localization patterns of the two proteins. Despite their overall differences in distribution, endogenous EB1 and Nav1 frequently co-localized at specific puncta. Some puncta were positive for both EB1 and Nav1, while other Nav1 puncta in close proximity showed no detectable EB1 (Figure 4C,D). Taken together, these data suggest that Nav1 is selectively targeted to structures within the T-zone, and are not exclusively found on growing microtubule ends.

### Differential Nav1 stoichiometry on growing microtubule ends in the growth cone

Specific targeting might enhance Nav1 association with the +TIP complex when polymerizing microtubules pass through the T-zone on their way to the P-domain. We therefore quantified the mean intensity of Nav1 and EB1 immunofluorescence at individual plus-ends, and we observed a nearly three-fold higher concentration of Nav1 on the P-domain localized plus-ends compared to the C-domain localized plus-ends (Figure 4E). In contrast, immunofluorescence intensity for EB1 on these same individual puncta did not differ. These data suggest that the stoichiometry of Nav1 interaction with plus ends can vary. They are consistent with the hypothesis that the T-zone serves as a reservoir for Nav1 and thereby enhances Nav1’s binding to the +TIP complex as microtubules polymerize toward the growth cone periphery, and suggest that microtubules could be one of the Nav1 effectors that regulate subcellular reorganization during growth cone morphogenesis.

### Nav1KO SH-SY5Y cells display transient membrane blebs that stem from dysregulation of the Rho/Rac pathway

Further observation of undifferentiated Nav1KO SH-SY5Y cells revealed that a portion of the cells displayed membrane blebs (Figure 5A). These membrane blebs were surrounded by a ring of F-actin and contained immunoreactivity for tubulin, reminiscent of non-apoptotic membrane blebs described elsewhere (Cunningham, 1995; Charras *et al*., 2008; Fackler and Grosse, 2008). Membrane blebs such as these occur when cortical F-actin disconnects from the plasma membrane, and the F-actin ring surrounding the blebs is thought to draw the plasma membrane back into the cell (Cunningham, 1995; Charras *et al*., 2008; Fackler and Grosse, 2008).

**Figure 5.**
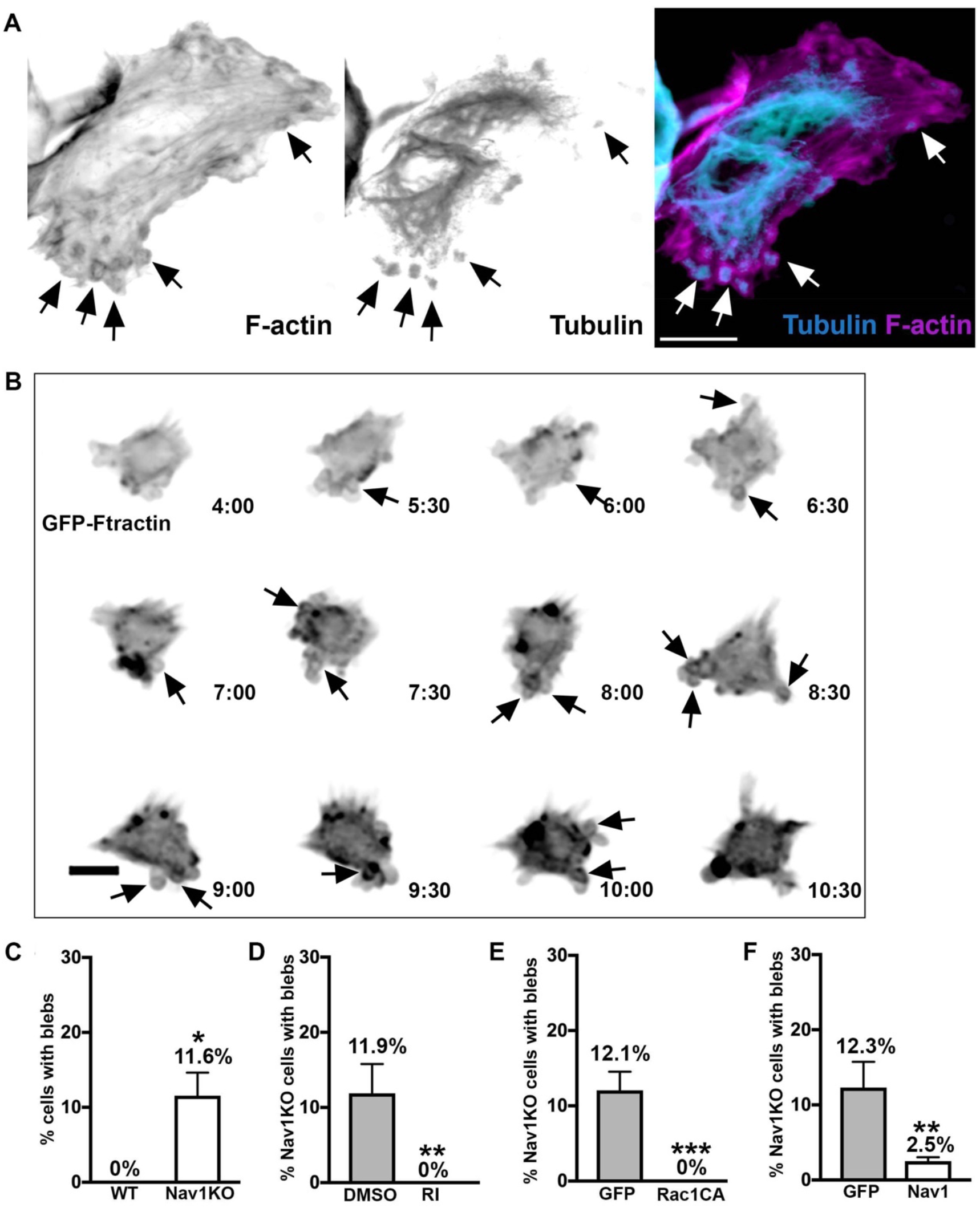
Nav1 KO cells reveal membrane blebs that indicate Nav1 regulates cortical actin dynamics via Rock and Rac1 pathway. (A) Representative image of SH-SY5Y Nav1KO cells displaying membrane blebs. Arrows indicate peripheral blebs. Scale bar = 10μm (B) Representative montage of a transiently blebbing cell expressing GFP-Ftractin for a portion of a 24 hour live imaging experiment. Cell imaged every 30 minutes, cell imaged every 30 minutes; time signature = hours:minutes. Scale bar = 10μm. (C) Nav1KO SH-SY5Y cells display significantly more membrane blebs than WT SH-SY5Y. Statistical analysis: Mann-Whitney, *p<0.05. n = 4 experiments (D) 2 hour ROCK Inhibitor (RI) treatment prevents blebbing in Nav1KO cells compared to Nav1KO cells treated with DMSO. Statistical analysis: Mann-Whitney,**p<0.01. n = 3 experiments (E) Nav1KO cells transfected with constitutively active Rac1 (Rac1CA) do not display membrane blebs compared to eGFP transfected Nav1KO cells. Statistical analysis: Mann-Whitney, ***p<0.001; n = 3 experiments (F) Electroporation of Nav1KO cells with GFP-Nav1 significantly reduced blebbing cells. Statistical analysis: Mann-Whitney, **p<0.01. n = 3 experiments All data are expressed as mean +/- SEM.

Observations of SH-SY5Y Nav1 KO cells via live imaging showed that blebs dynamically extended and retracted from the plasma membrane, and that periods of membrane blebbing were transient in any given cell. Time-lapse imaging over 24 hours indicated that, on average, 51% of Nav1KO cells displayed transient membrane blebs for periods of time (Fig 5B, black arrows point to blebs). In contrast, blebbing cells were never observed in the WT line. In fixed samples we found that, on average, 12 percent of Nav1KO cells displayed membrane blebs (Figure 5C). This suggests that the majority of cells undergo membrane blebs at some point over the course of time, and that fixed cells capture only a small portion of blebbing cells at any given moment. Together, these data suggest that Nav1 may be important for the proper connection of cortical actin and the plasma membrane, which is a previously undescribed role for Nav1.

A balance in the activities of RhoA and Rac1 is important for maintaining the cell cortex. Specifically, over-active RhoA or under-active Rac1 can lead to membrane blebbing (Chauhan *et al*., 2011; Bang *et al*., 2015). Therefore, we investigated the role of RhoA and Rac1 in the Nav1KO membrane blebs. To test the role of RhoA, we incubated Nav1KO cells with the Rho-kinase (a primary RhoA effector molecule) inhibitor Y-27632 (5µM), and found it completely rescued the membrane blebs compared to DMSO-treated Nav1KO cells (Figure 5D). To test Rac1, we expressed an eGFP control or a constitutively active form of Rac1 (Rac1CA) in Nav1KO cells, and found that Rac1CA also completely rescued the membrane bleb phenotype (Figure 5E). To ensure that the membrane blebs are attributable to the loss of Nav1 we performed a rescue experiment. Reintroduction of GFP-Nav1 to Nav1KO cells significantly decreased the number of cells with membrane blebs (Figure 5F). These results collectively suggest that Nav1 promotes a proper balance in the Rho/Rac pathway, and that a Nav1 deficit results in overactive Rho-kinase and/or under-active Rac1, resulting in membrane blebs.

### Nav1 accumulates in and promotes F-actin rich membrane ruffles

Because we observed that a substantial portion of Nav1 puncta in the growth cone are not resident on growing microtubule ends and that Nav1 plays a role in actin organization in the cell periphery, we investigated the localization of Nav1 puncta in relation to actin-rich growth cone structures in hippocampal neurons. Immunoreactivity for endogenous Nav1 displayed a highly punctate pattern within all three domains of the growth cone (the C-domain, P-domain, and T-zone). However, Nav1 clusters of various sizes and shapes were usually enriched within the T-zone (Figure 6A,B), a region that lies just distal to the microtubule-filled C-domain (Figure 6A) and is enriched in cortactin (Figure 6B) and drebrin, actin binding proteins concentrated in active membrane ruffles (Weed *et al*., 1998; Shirao *et al*., 2017). Ectopic GFP-Nav1 strongly co- localized with phalloidin labeled F-actin in sub-domains within the growth cone (Figure 6C).

**Figure 6.**
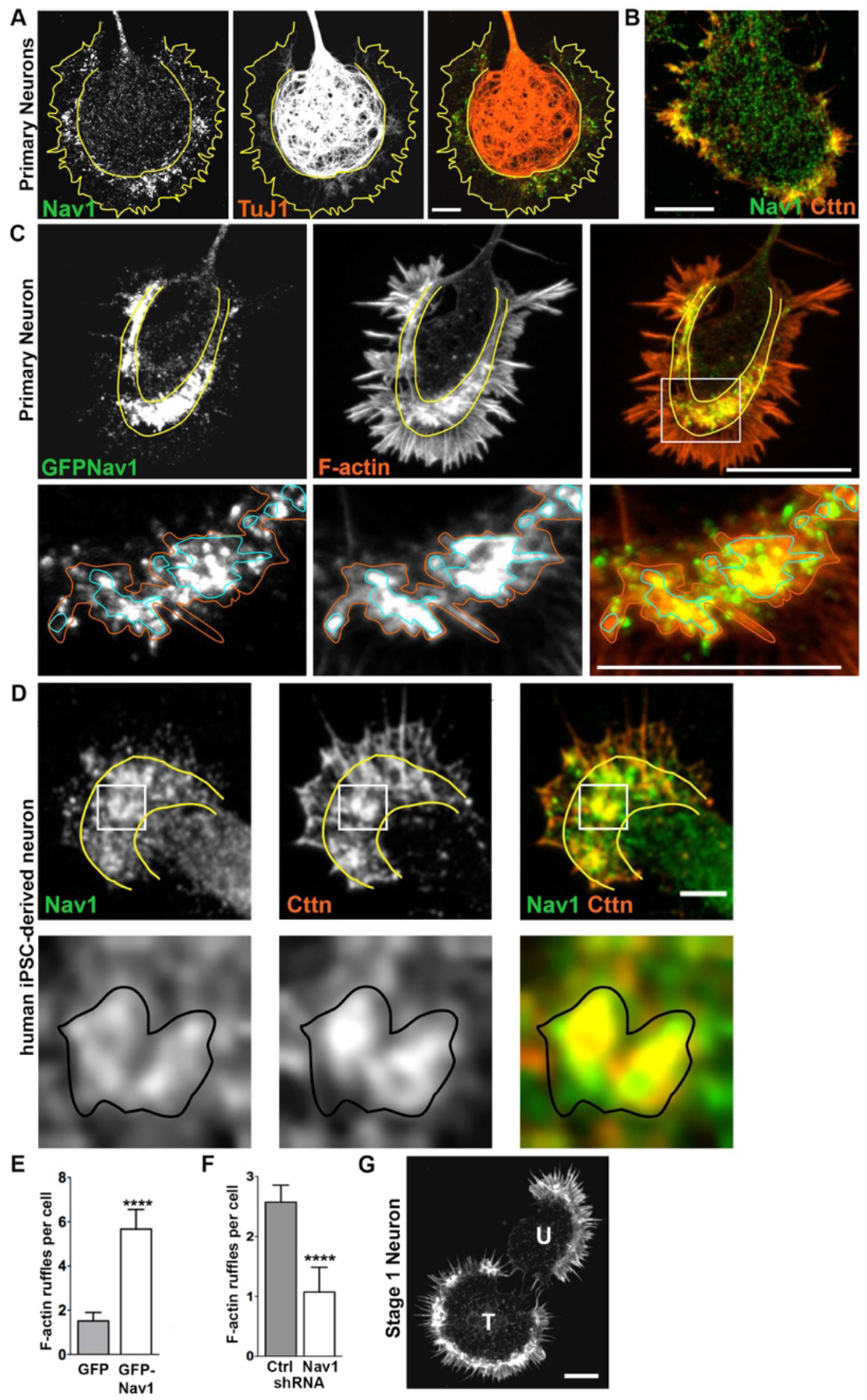
Nav1 associates with and regulates F-actin membrane ruffles in the transition zone in the growth cone. (A) Growth cone stained for endogenous Nav1 and the actin binding protein cortactin (Cttn), which is a marker of actin-rich membrane ruffles. Scale bar = 20μm (B) Cultured rat embryonic neurons were fixed on 3DIV and double labeled for endogenous Nav1 and ß-III tubulin (TuJ1). Growth cone domains borders are highlighted in solid yellow lines; P, peripheral domain; T, transition zone; C, central domain. Note the accumulation of endogenous Nav1 puncta within clusters in the T zone. Scale bar = 10μm (C) Growth cone from a stage 2 hippocampal neuron expressing GFP-Nav1, labeled for F-actin by Alex568-conjugated phalloidin. The three principal growth cone subdomains are labeled as in (a). Boxed region is enlarged in panel directly below. Note that clusters of GFP-Nav1 puncta are enriched within the T-zone, and their concentration correlates with that of F-actin. Orange and blue solid lines indicate regions of moderate and intense concentrations, respectively, of F-actin, which correspond to membrane ruffles. Scale bars: upper panel 20μm, lower panel, 10μm (D) Differentiated human iPSC-derived neurons were fixed and double labeled for endogenous Nav1 and Cttn. Boxed region is enlarged in panel directly below. Black solid lines indicate membrane ruffle where both Cttn and Nav1 are enriched. Scale bar = 10μm. All data are expressed as mean +/- SEM (E) GFP- Nav1 over-expression increases the number of F-actin ruffles per neuron. Mann-Whitney test, ****p<0.0001. n = 3 experiments; GFP transfected = 29 cells; GFP-Nav1 transfected = 30 cells. (F) Knockdown of endogenous Nav1 using shRNA decreases the number of F-actin rich membrane ruffles per neuron. Statistical analysis same as (D). Control and Nav1 shRNA = 70 cells each (G) Two stage 1 neurons stained with Alexa568- conjugated phalloidin. Note that the cell transfected with GFP-Nav1 (“T”) shows large accumulations of F-actin within the transition zone compared to the adjacent untransfected neuron (”U”). Scale bar = 20μm. All data are expressed as mean +/- SEM.

With higher magnification we observed that, although there was not a perfect spatial overlap between GFP-Nav1 and phalloidin-labeled F-actin, clearly the most intense accumulations of GFP-Nav1 puncta corresponded to regions where phalloidin staining was similarly most intense (Figure 6C, see zoomed region depicted in lower panels). In neurons co-transfected with GFP- Nav1 and mRFP1-actin, we observed that such regions dynamically changed size, shape, and location in a coordinated manner. In dynamic behavior these Nav1 and F-actin enriched regions were reminiscent of actin–rich membrane ruffles (Supp. Figure 4). We therefore conclude that Nav1 is highly enriched in membrane ruffles, and use this term to refer to these structures throughout this study.

Growth cones in SH-SY5Y cells are typically small, and we did not succeed in finding culture conditions to facilitate the development of large, well-spread growth cones. However, in larger size growth cones in human iPSC-derived neurons where a T-zone could be discerned we observed clusters of Nav1 puncta enriched in the T-zone of the growth cone colocalized with cortactin (Figure 6D), suggesting that T-zone accumulation of Nav1 puncta is a general feature.

Strikingly, Nav1 not only accumulated in membrane ruffles in the T-zone but also regulated their presence. Expression of GFP-Nav1 in hippocampal neurons (which on average resulted in ∼2.5- fold increase over endogenous levels) induced a significant increase in the number of actin-rich ruffles per cell (Figure 6E). Endogenous Nav1 similarly regulates membrane ruffles, since shRNA against Nav1 induced a significant decrease in membrane ruffles compared to control neurons (Figure 6F). Interestingly, we also observed that stage 1 neurons (i.e., pre-neurite initiation) that expressed GFP-Nav1 displayed enriched F-actin in the lamellipodial region in membrane ruffles (Figure 6G), suggesting that lamellipodial regions from which growth cones presumably emerge also display these Nav1-associated structures. Collectively, these data indicate that Nav1 has a concentration-dependent effect on the presence of membrane ruffles in the developing growth cone.

### Nav1-rich membrane ruffles are sites of bulk endocytosis

While growth cone membrane ruffles and similar dynamic actin-rich structures have been described previously, (Thompson *et al*., 1996; Rochlin *et al*., 1999; Bonanomi *et al*., 2008; Buck *et al*., 2017), their specific function at the growth cone is not well-defined. One function proposed for growth cone membrane ruffles in the early growth cone is non-clathrin mediated bulk endocytosis (Diefenbach *et al*., 1999; Bonanomi *et al*., 2008). We therefore used the lipophilic dye FM4-64 to examine endocytosis at the growth cone (Bonanomi *et al*., 2008). This dye is readily endocytosed in a non-specific manner after attaching to the membrane (Betz *et al*., 1996). As expected, we observed that FM4-64 is taken up in areas of the growth cone with F-actin membrane ruffles. To test whether enriched clusters of Nav1 puncta in the growth cone are sites of bulk endocytosis, we electroporated GFP-Nav1 and pcs-CeruleanMembrane-FP (to visualize the whole growth cone) into rat primary neurons, and repeated the FM4-64 experiments. Indeed, we found that FM4-64 uptake at the growth cone occurs where GFP-Nav1 and F-actin are enriched (Figure 7A). Furthermore, we compared growth cones expressing GFP- Nav1 and pcs-CeruleanMembrane-FP to growth cones expressing pcs-CeruleanMembrane-FP alone, and found that growth cones expressing ectopic Nav1 showed significantly more FM4-64 uptake (Figure 7B).

**Figure 7.**
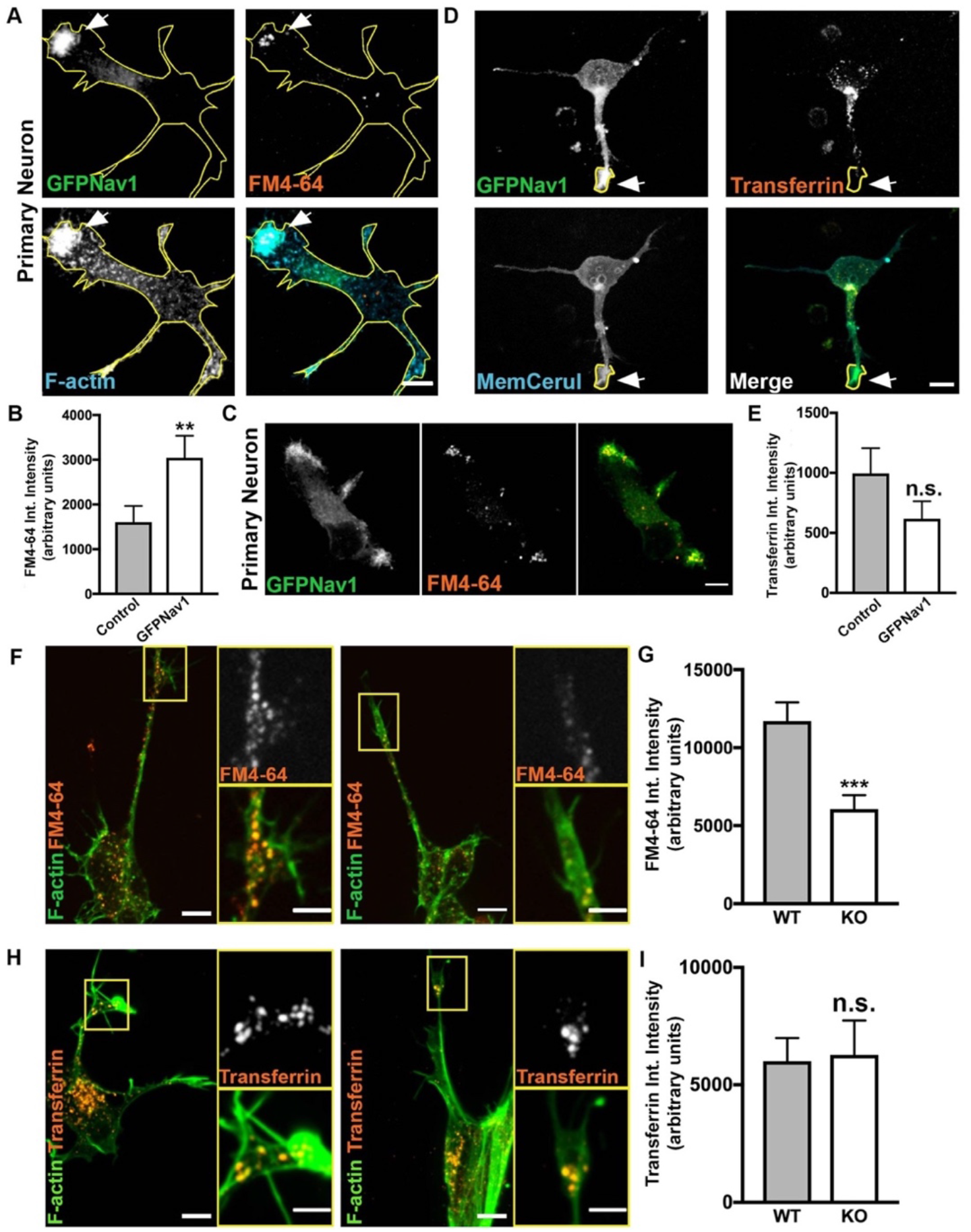
Nav1 promotes endocytosis at F-actin membrane ruffles. (A) Representative image of 3DIV hippocampal neurons transfected with GFP-Nav1 (green) pcs-CeruleanMembrane-FP (used to make yellow cell outline), incubated with FM4-64 (red), and incubated with Alexa647 conjugated Phalloidin (blue) demonstrates Nav1-enriched F-actin ruffles are sites of FM4-64 uptake. Arrow points to region enriched in GFP-Nav1, F-actin and FM4-64. (B) Growth cones expressing GFP-Nav1 and pcs- CeruleanMembrane-FP have significantly more FM4-64 uptake than growth cones just expressing pcs-CeruleanMembrane-FP. Statistical analysis: Mann-Whitney, **p<0.01. n = 4 experiments; Control = 69 growth cones GFP-Nav1 = 69 growth cones (C) Representative image of 3DIV stage 1 hippocampal neuron transfected with GFP-Nav1 (green) and incubated with FM4-64 (red) demonstrating that FM4-64 Scale bar = 10μm (D) Representative image of 3DIV hippocampal neurons transfected with GFP- Nav1 (green) and pcs-CeruleanMembrane-FP (blue) and incubated with 568-conjugated transferrin (red). Scale bar = 10μm (E) There is no difference in transferrin intensity between GFP-Nav1 expressing and control growth cones. Statistical Analysis: Mann-Whitney test, n.s. = not significant. n = 4 experiments; Control = 34 growth cones, GFP-Nav1 = 42 growth cones (F) Representative image of 7 day differentiated WT and Nav1KO SH-SY5Y cells with phalloidin-labeled F-actin (green) and incubated with FM4-64 (red). Scale bar = 10μm, 5μm for zoomed image. (G) Nav1KO SH-SY5Y growth cones have less FM4- 64 uptake than WT cells, as measured by FM4-64 intensity. Statistical analysis: Mann-Whitney, ***p<0.001. n = 4 experiments; WT = 185 growth cones, KO = 116 growth cones. (H) Representative images of 7 day differentiated WT and Nav1KO SH-SY5Y cells with phalloidin-labeled F-actin (red) and incubated with transferrin (red). Scale bar = 10μm, 5μm for zoomed image. (I) There is no significant difference in transferrin intensity between WT and Nav1KO SH-SY5Y cells. Statistical analysis: Mann- Whitney, n.s. = not significant. n = 4 experiments; WT = 83 growth cones, KO = 59 growth cones. All data are expressed as mean +/- SEM.

When we observed stage 1 neurons, we found that they, too, had FM4-64 uptake at Nav1 enriched peripheral membrane ruffles (Figure 7C), suggesting that Nav1-promoted endocytosis occurs from the earliest stages of neuronal development. We also found that GFP-Nav1 expressing growth cones displayed significantly higher intensity of pcs-CeruleanMembrane-FP, suggesting that Nav1 may promote membrane accumulation (Supp. Fig 5A,B). These data suggest a novel role of Nav1 in regulating bulk endocytosis and membrane recycling at the growth cone.

To test whether Nav1 promotion of endocytosis is specific to non-clathrin mediated endocytosis, we tested uptake of transferrin in primary neurons, as transferrin is taken up exclusively by clathrin-mediated endocytosis (Hinrichsen *et al*., 2003). We found no difference in transferrin uptake in GFP-Nav1 expressing neurons compared to control, (Figure 7D,E). Furthermore, there were substantially fewer GFP-Nav1-transfected or untransfected growth cones with any detectable transferrin uptake compared to growth cones exhibiting FM4-64 uptake (GFP-Nav1 transfected growth cones: 60% FM4-64 uptake, 26% transferrin uptake; untransfected growth cones: 40% FM4-64 uptake,14% transferrin uptake). Our results are consistent with the previous conclusion that clathrin-mediated endocytosis is not a predominant means of endocytosis in the neuronal growth cone during early morphogenic stages (Bonanomi *et al*., 2008).

We also confirmed that endocytosis occurs in F-actin rich and GFP-Nav1 enriched membrane ruffles in differentiated SH-SY5Y cell growth cones (Supp. Figure 5C), and found that a significantly lower amount of FM4-64 was taken up in growth cones in the Nav1KO cells compared to WT controls (Figure 7F,G). This directly implicates endogenous Nav1 in endocytosis at the growth cone. We also compared transferrin uptake in growth cones of WT versus Nav1KO SH-SY5Y cells, and found no difference between the two lines (Figure 7H,I). Altogether, our Nav1 gene silencing and overexpression data demonstrate that Nav1 specifically promotes non-clathrin-mediated endocytosis in growth cones.

### Nav1 promotes macropinocytosis

We hypothesized that the Nav1-regulated endocytosis occurring at the growth cone is the same form of micropinocytosis identified by Bonanomi *et al* (2008). To test this hypothesis, we first ectopically expressed GFP-Nav1 in primary neurons, and incubated them with high molecular weight (70kDa) dextran, uptake of which is a marker of macropinocytosis (Falcone *et al*., 2006). We found that 70kDa dextran is indeed taken up in Nav1-enriched growth cones membrane ruffles (Figure 8A). Next, we blocked some reported molecular components of macropinocytosis to test whether that would decrease GFP-Nav1-mediated FM4-64 uptake. First, we used methyl- β-cyclodextrin (MβCD) to deplete cholesterol and found that this treatment significantly decreased FM4-64 uptake in GFP-Nav1-expressing neurons (Figure 8B). Macropinocytosis is also reliant on phosphoinositide-3 kinase (PI3K) (Araki *et al*., 1996; Rupper *et al*., 2001), and indeed we observed that the PI3K inhibitor LY294002 significantly decreased the percentage of growth cones that had FM4-64 uptake (Figure 8C). Together, the uptake of 70kDa dextran at GFP-Nav1-enriched growth cones, along with the requirement of cholesterol and PI3K for FM4- 64 uptake in the presence of GFP-Nav1 strongly supports the conclusion that Nav1 promotes a macropinocytosis form of bulk internalization at the growth cone.

**Figure 8.**
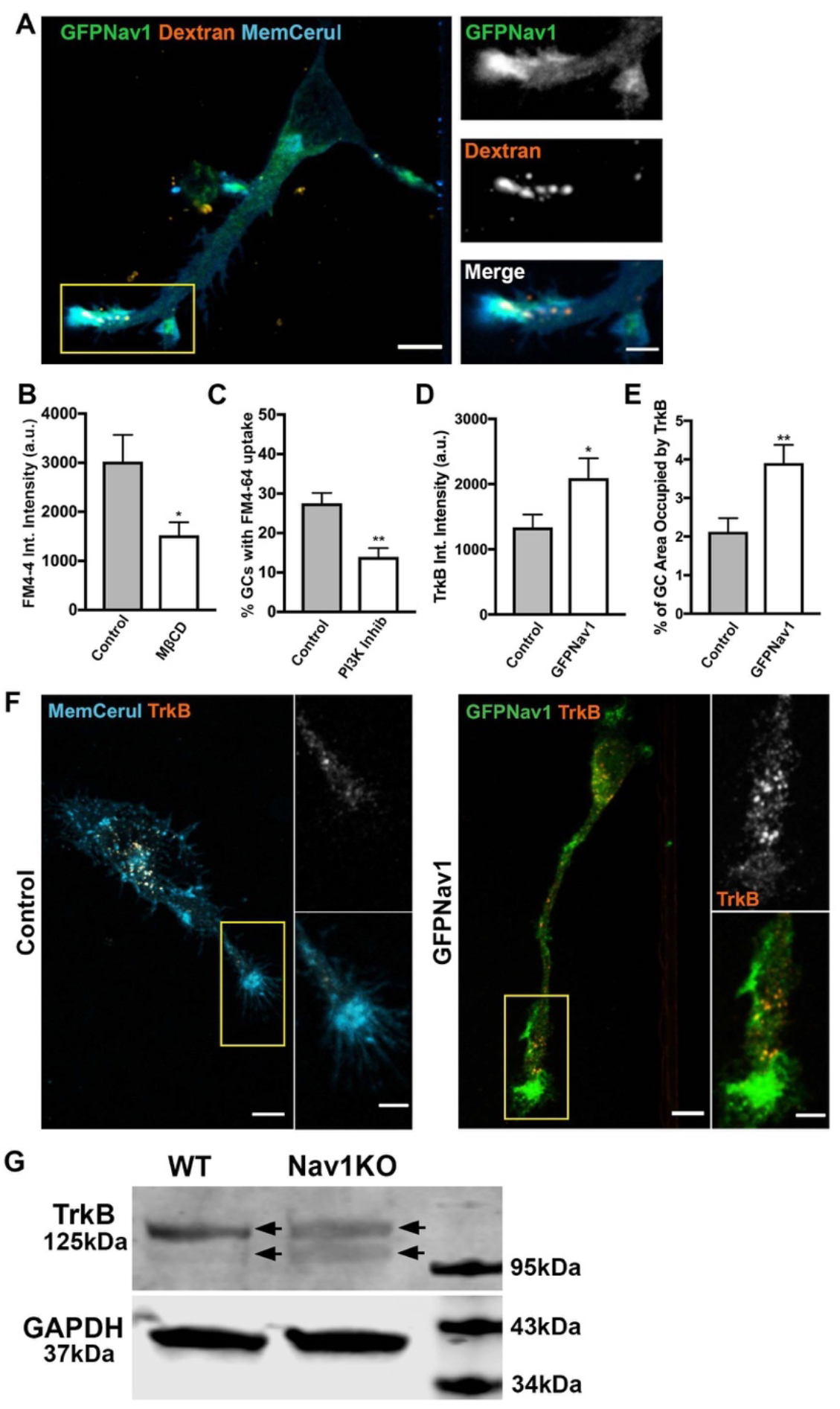
Nav1 promotes macropinocytosis and growth factor internalization at the growth cone. (A) Representative image of 3DIV hippocampal neurons transfected with GFP-Nav1 (green) and pcs-CeruleanMembrane-FP (blue) and incubated with 70kDa Dextran (red) showing high molecular weight dextran is taken up in Nav1-enriched growth cones. Scale bar = 10μm and 5μm for zoomed images. (B) MßCD treatment of GFP-Nav1 transfected primary neurons decreases FM4-64 uptake in growth cones. Statistical analysis: Mann-Whitney, **p<0.01. n = 3 experiments; Water = 46 growth cones, MßCD = 53 growth cones. (C) PI3 kinase inhibition of GFP-Nav1 transfected primary neurons decreases the number of growth cones with FM4-64 uptake. Statistical analysis: Mann-Whitney, *p<0.05, n = 4 experiments. (D) Growth cones of primary neurons expressing GFP-Nav1 have significantly more TrkB internalization by intensity than control growth cones. Statistical analysis: Mann-Whitney, *p<0.05. n = 3 experiments; Control = 61 growth cones, GFP-Nav1 = 60 growth cones (E) Growth cones of primary neurons expressing GFP-Nav1 have significantly more TrkB internalization by percent area occupied than control growth cones. Statistical analysis: Mann-Whitney, ***p<0.001, n same as (D). (F) Representative images of control and GFP-Nav1 primary neurons, expressing pcs-CeruleanMembrane-FP or GFP-Nav1 and pcs-CeruleanMembrane-FP, respectively, after internalization of the TrkB 1D7 antibody. Scale bar = 10μm, 5μm for zoomed images. (G) Representative western blot of 4 day retinoic acid differentiated WT and Nav1KO SH-SY5Y cells showing a double band (arrows indicate each band of doublet) for TrkB. GAPDH included as a loading control. All data are expressed as mean +/- SEM.

### Nav1 regulates internalization of TrkB in growth cones

The function of endocytosis within growth cone membrane ruffles remains poorly characterized. Previous studies showed that Nav1 is critical for the neurite outgrowth response to the guidance cue netrin (Martinez-Lopez *et al*., 2005; Sanchez-Huertas *et al*., 2020). Netrin is one of many guidance cues that direct axon outgrowth and neuronal migration via signaling to the cytoskeleton. Such cues activate a variety of receptors, and some are internalized via endocytosis while others are not (Winckler and Yap, 2011). TrkB is a receptor for the growth factor BDNF, is taken up to influence cell growth and differentiation, and can be internalized via macropinocytosis (Valdez *et al*., 2005; Winckler and Yap, 2011; Scott-Solomon and Kuruvilla, 2018). Because of this, we hypothesized that Nav1 might promote internalization of the BDNF receptor TrkB, which provides critical signals for neuronal morphogenesis and function. To test this hypothesis, we used the 1D7 TrkB antibody, which can be applied to live cells, internalized, then visualized via imaging (Bai *et al*., 2010). We found that ectopic expression of GFP-Nav1 lead to a significant increase in TrkB uptake at hippocampal growth cones (Figure 8D-F). These data demonstrate that Nav1 promotes endocytosis of growth factors important for neuritogenesis.

To test whether this internalization difference could be due to altered expression of TrkB, we performed western blots in differentiated SH-SY5Y cells. We found no quantitative difference in TrkB expression levels, but, interestingly, we observed that while TrkB immunoreactive signal appeared as a doublet in lysates from both WT and Nav1KO cells, the lower band from the WT cells was notably fainter than that in the Nav1KO cells (Figure 8G).

## DISCUSSION

### Nav1 is enriched in pathfinding axons and areas of morphological change to promote neuritogenesis and growth cone formation

Previous research and the data presented in this paper demonstrate that Nav1 is a cytoskeleton-associated protein, able to influence actin dynamics in multiple settings, including neural development (Maes *et al*., 2002; van Haren *et al*., 2009; van Haren *et al*., 2014; Sanchez-Huertas *et al*., 2020). Dysregulation of the cytoskeleton is implicated as an underlying mechanism associated with many neurodevelopmental disorders, including lissencephalies, ASD, and intellectual disability (des Portes *et al*., 1998; Bellenchi *et al*., 2007; Bamba *et al*., 2016; Kannan *et al*., 2017), underscoring the complex and precise cytoskeletal regulation necessary for early brain development.

Here we show that Nav1 is enriched in areas where rapid cell morphogenesis occurs, including pathfinding axons *in vivo* as well as in the cell periphery and distal neurites in multiple neural cell types: primary rodent neurons, differentiated SH-SY5Y cells, and human iPSC-derived neurons. Additionally, we show that reduction of Nav1 via shRNA-knockdown or complete knockout inhibits neuritogenesis in primary neurons and SH-SY5Y cells, respectively. Defective neuritogenesis after Nav1 knockdown was demonstrated previously in neuroblastoma cells (van Haren *et al*., 2014), and defective neurite outgrowth in response to netrin was also shown (Martinez-Lopez *et al*., 2005; Sanchez-Huertas *et al*., 2020). However, our study is the first to show that neurite initiation, in addition to elongation, is affected in neurons with Nav1 depletion. We also provide evidence for altered morphology and guidance after Nav1 knockdown *in vivo*. A growing body of evidence thus implicates an integral role for Nav1 in the cell’s ability to organize the cytoskeleton to properly produce, elongate, and guide functional neurites during differentiation.

### Nav1 promotes cortical actin-membrane association via Rho-GTPase signaling

We demonstrated that Nav1-knockdown induces non-apoptotic transient membrane blebbing that denotes a disruption in the cortical actin-membrane association. Membrane blebs in the Nav1KO cells were rescued by either Rho-kinase inhibitor or introduction of a constitutively active Rac1, suggesting that disrupted Rac1 and Rho-kinase signaling contribute to membrane blebbing in the Nav1KO cells. Nav1 has been shown to promote Rac1 activation in the context of neuritogenesis via the guanine nucleotide exchange factor (GEF) Trio (van Haren *et al*., 2014), a gene that has been implicated in autism (Sadybekov *et al*., 2017). It is therefore logical that a total knockout of Nav1 would disrupt Rac1 signaling. Furthermore, Rac1 and Rho-kinase often have opposite effects on cytoskeleton dynamics in a variety of cellular processes (Da Silva *et al*., 2003; Symons and Rusk, 2003; Chauhan *et al*., 2011; Hodge and Ridley, 2016), and disruption of Rho signaling has been shown to cause membrane blebs in other cell types (Wilkinson *et al*., 2005; Fackler and Grosse, 2008; Sanz-Moreno *et al*., 2008; Honnappa *et al*., 2009). These data suggest that Nav1 may have a larger influence on Rho signaling than previously known, and suggest a novel connection among Nav1, the cytoskeleton, and regulation of the plasma membrane.

### Nav1 is associated with +TIPs in the growth cone

In accordance with previous studies, we confirm that Nav1 tracks growing microtubule ends in association with EB in the growth cone (Martinez-Lopez *et al*., 2005; van Haren *et al*., 2009; van Haren *et al*., 2014; Sanchez-Huertas *et al*., 2020). Importantly, we observe a significantly higher concentration of Nav1 on individual microtubule plus ends in the P-domain versus the C-domain. This suggests that Nav1 stoichiometry can potentially vary on microtubule plus ends, an observation that warrants further study. Microtubules and +TIP proteins can aid in delivery of necessary molecules for growth factor response signaling to the growth cone periphery, and can also influence a diversity of cellular processes (Williamson *et al*., 1996; Akhmanova and Steinmetz, 2008; Lowery and Van Vactor, 2009). For example, neurotrophin receptors are incorporated into early endosomes in the growth cone and then retrogradely transported on microtubules (Delcroix *et al*., 2003). Thus, we speculate that the peripheral Nav1-occupied microtubule plus ends may represent a subset of +TIPs poised for endosomal transport at sites of uptake.

### Nav1 accumulates in and regulates T-zone F-actin rich membrane ruffles and endocytosis

The transition zone of the growth cone is of potential importance for multiple functions for growth cone guidance. The T-zone may act as a microtubule barrier, permitting only a select set of pioneering microtubules to advance to the growth cone periphery (Medeiros *et al*., 2006; Bearce *et al*., 2015). This “gatekeeper” function may represent a key regulator of signaling to influence growth cone morphological changes (Medeiros *et al*., 2006; Bearce *et al*., 2015). The T-zone is also a site of membrane recycling, ensuring that membrane is available for dynamic changes in the growth cone (Diefenbach *et al*., 1999; Bonanomi *et al*., 2008). We observed that clusters of Nav1 puncta are enriched specifically in the actin-rich regions of the T-zone, and that these actin-rich accumulations resemble membrane ruffles. Furthermore, we demonstrated that Nav1 promotes actin ruffles in the T-zone, indicating that Nav1 may promote the formation or persistence of the ruffles.

The function of growth cone membrane ruffles is relatively uncharacterized, but in non-neuronal cell types, membrane ruffles are associated with actin-remodeling and cell migration (Mitchison and Cramer, 1996; Innocenti, 2018; Schnoor *et al*., 2018). Prior evidence suggests that growth-cone membrane ruffles are sites of bulk endocytosis, and that the endocytosis is necessary for neurite outgrowth (Bonanomi *et al*., 2008). We observed patches of enriched F-actin as sites of non-clathrin mediated, but not clathrin-mediated endocytosis in our rat primary neuron cultures, and we demonstrated that Nav1 promotes endocytosis in in these regions. This is a novel mechanism by which Nav1 may influence neuritogenesis. There are several mechanisms by which Nav1 might promote endocytosis, and this presents an opportunity for further research.

For example, as described above, Nav1 promotes Rac1 activation through its activation of the GEF Trio (van Haren *et al*., 2014), and Rac1 promotes membrane ruffles (Ridley *et al*., 1992), as well as bulk endocytosis (Symons and Rusk, 2003; Bonanomi *et al*., 2008). Moreover, Rac1, along with other Rho-family proteins, plays an important role in endocytosis regulation.

Therefore, we postulate that one key function of Nav1 is to influence non-clathrin mediated endocytosis via its interaction with Trio. It is also possible that Nav1 influences endocytosis by promoting actin polymerization at the membrane, which in turn promotes endocytosis. A recent study reported that Nav1 and actin may interact directly via the +TIP binding domain (Sanchez-Huertas *et al*., 2020). Thus, Nav1 might stimulate endocytosis through direct binding to F-actin as an additional or an alternative means to its regulation of Rac1.

Other dynamic actin-rich structures have been described in the growth cone literature. Inductapodia are dynamic actin-rich structures that form in *Aplysia* growth cones as localized sites where the growth cone plasma membrane engages integrin-dependent cell adhesion molecules (Thompson *et al*., 1996; Rochlin *et al*., 1999; Buck *et al*., 2017), and their formation is dependent upon Rac1 activity (Buck *et al*., 2017). Intrapodia are described in rodent sensory neurons as actin-rich structures of unknown function that form spontaneously in the growth cone membrane at microtubule tips (Rochlin *et al*., 1999). Further research is required to determine what relationship, if any, such structures have to the actin-rich membrane ruffles described here and in Bonanomi *et al* (Bonanomi *et al*., 2008).

The function of membrane ruffle macropinocytosis is not well understood, but our data suggest a novel connection to neurotrophin signaling. We found that ectopic expression of Nav1 significantly increased the uptake of TrkB, the receptor for the growth and guidance cue BDNF, in the growth cone. TrkB is endocytosed via both clathrin mediated endocytosis (Beattie *et al*., 2000; Zheng *et al*., 2008), and clathrin-independent macropinocytosis (Valdez *et al*., 2005; Xu *et al*., 2016). Our study implicates Nav1 in non-clathrin mediated macropinocytic TrkB uptake as a mechanism by which Nav1 promotes neuronal responses to growth and guidance cues. Indeed, we found that SH-SY5Y Nav1 KO cells were defective in BDNF-stimulated neuritogenesis.

Furthermore, our immunoblot results suggest that Nav1 gene silencing causes an alteration in the molecular properties of the BDNF receptor TrkB. The observed alteration in the relative mobility of TrkB in SDS-PAGE suggests that one or more transcriptional, translational, or post-translational modifications is somehow under Nav1’s control, which is an interesting topic for further investigation. As a receptor tyrosine kinase, TrkB becomes autophosphorylated on multiple tyrosine residues upon ligand engagement and activation (Middlemas *et al*., 1994), a step that is essential for recruitment of downstream effectors to mediate cell signaling for neuronal survival and growth (Kaplan and Miller, 2000). Additionally, TrkB is phosphorylated by Cdk5 on Serine478, which is essential for BDNF promotion of dendritic growth and long-term potentiation of synaptic plasticity (Cheung *et al*., 2007; Lai *et al*., 2012). Phosphorylated TrkB receptor-mediated intracellular signaling is integral to neuronal survival and neurite outgrowth (Scott-Solomon and Kuruvilla, 2018), and this therefore represents a potential downstream role for Nav1. Furthermore, it has been shown that endocytosis of TrkB receptors is necessary for directional migration of cerebellar granule cell precursors in response to BDNF (Zhou *et al*., 2007). In this manner, disruption of Nav1-mediated TrkB endocytosis may explain the leading-process misorientation phenotype we observed upon Nav1 reduction in developing mouse cortex (Figure 3H,J). The importance of tight regulation of neurotrophin endocytosis for neural morphogenesis during brain development and the role of Nav1 in these processes opens up an exciting avenue of research.

## Conclusions

Our data suggest a novel concept regarding the function of Nav1 in the regulation of neuronal development. Nav1 not only associates selectively with and regulates both the microtubule and actin cytoskeletons, but may play a general role in mediating uptake of and signaling by neurotrophins and other growth and guidance cues. This is especially important in developing neurons, where Nav1 is highly expressed, and allows newly extending axons to quickly respond to extra- and intracellular signals to facilitate tightly-regulated morphogenesis and brain circuit establishment.

## METHODS

### Reagents

Pharmacological compounds were obtained from the following sources: Rho Kinase Inhibitor (Tocris). Retinoic Acid (Sigma), Brain-derived neurotrophic factor (BDNF) (Peprotech), Methyl-*β*-Cyclodextrin (M*β*CD) (Sigma), phosphoinositide-3 kinase (PI3Kinase) inhibitor LY294002 (Selleck Chemicals).

### Plasmids

mRFP-*β*-actin (Calabrese and Halpain, 2005), GFP-Ftractin (kind gift from Higgs lab), GFP-Nav1 (Martinez-Lopez *et al*., 2005), mVenus (Hand and Polleux, 2011), pECFP-RacQ61L (kind gift from Dr. K. Yamada, Addgene #105292), pEGFP-N1 (Clontech), pCS-membrane-ceruleanFP (kind gift from Dr. S. Megason, Addgene #53749), EB3mCherry (kind gift from M. W. Davidson). Nav1 shRNA #1 and #2 were inserted into the pSuper vector according to manufacturer’s instructions (Oligoengine) (Supp. Table1). However, shRNA#1 outperformed shRNA#2 in knocking down exogenous Nav1 in neurons (Supp. Fig2A).

### Cell Culture and Differentiation

All cell lines and primary neurons were cultured at 37°C and 5% CO_2_. N1E-115 cells were cultured in Dulbeco’s modified Eagles medium (DMEM; Gibco) supplemented with 10% (v/v) fetal bovine serum (FBS; Atlanta Biologicals S11150). SH-SY5Y (ATCC CRL-2266) cells were cultured in DMEM/F12 1:1 with 25mM HEPES, 10% heat inactivated FBS, and penicillin/streptomycin. The SH-SY5Y differentiation protocol was modified from Encinas *et al (2000)*. Once undifferentiated SH-SY5Y cells reached 90% confluency, they were rinsed once with PBS, and then RA media (DMEM/F12 with 0.5% FBS, and 10µM Retinoic Acid) was added. RA media was changed on day 2 of differentiation, and were replated on day 4 using StemPro accutase (Life Technologies); cells were seeded at 394 cells/mm^2^ on ibidi µ-Plate 24 well (82406) coated with 200µg/mL poly-(D)-lysine overnight at room temperature. Cells were cultured in RA media for 24 hours after re-plating, and then media was changed to BDNF media (DMEM/F12 and 50ng/µL BDNF), with half media changes every other day. Nucleofected SH-SY5Y cells were plated into 6 well plates dishes for 24 hours to recover, and then cells were seeded at 394 cells/mm^2^ on ibidi µ-Plate 24 well coated with 100µg/mL poly-(D)-lysine, and fixed 24 hours later.

Human iPSCs were purchased from Cellular Dynamics (iCell GlutaNeurons R1061), and cultured per the manufacturer’s protocol. Primary hippocampal cultures were prepared from embryonic day 19 rat brains as described (Calabrese and Halpain, 2014). For fixed primary neuron experiments (except FM4-64 experiments), cells were plated on coverslips coated with poly-(D)-lysine (Sigma 100ug/mL) at a density of 375 cells/mm^2^.

### Transfection methods

### Dissociated primary neurons and undifferentiated SH-SY5Y cells were electroporated, prior to plating, using either Lonza Nucleofector 2b, or Lonza 4D-Nucleofector core unit, according to manufacturer’s protocol

#### Lipofection of cultured cells

N1E-115 cells and SH-SY5Y cells (Rac1Q61L experiment and membrane blebs live imaging) were transfected using either Lipofectamine 2000 or 3000 (ThermoFisher Scientific), respectively, according to the manufacturer’s protocol. Nav1 shRNA and control vector was incubated for 3 days, all other constructs were incubated for 24 hrs.

#### In utero electroporation

Electroporation was performed as previously described (Courchet *et al*., 2013). Endotoxin-free plasmid DNA was injected at a concentration of 1 μg/μl in the lateral ventricles. mVenus was co-electroporated at a 2:1 ratio to the pSuper empty vector or Nav1 shRNA to verify that fluorescent neurons were electroporated with the DNA of interest. Electroporation was performed with gold-coated paddles at E15.5 to target cortical layer 2/3. The electroporation settings were: 4 pulses of 45V for 50 ms with 500 ms interval.

### Nav1 antibody generation

Anti-Nav1 was produced using the GST-MTD plasmid construct expressed in *E.coli*. Soluble GST-MTD was purified on a glutathione affinity column, eluted with glutathione, and injected into rabbits (antiserum production by COVANCE). Antisera to GST-MTD was affinity purified by pre-absorption on an Affi-Gel (Bio-Rad) column coupled with GST, followed by affinity purification with an Affi-Gel column coupled with GST-MTD. This affinity-purified antibody was used for all experiments to detect endogenous Nav1. Antibody specificity to Nav1 was confirmed as follows: using immunoblot, lack of detectable immunoreactivity for the Nav1 antibody in extracts of Nav1 KO SH-SY5Y cells compared to WT controls (Suppl. Fig. 2B); using immunofluorescence staining, lack of detectable immunoreactivity for the Nav1 antibody in of Nav1 KO SH-SY5Y cells compared to WT controls (Suppl. Fig. 2C).

### SH-SY5Y Nav1KO cell line generation

Nav1 Knock-out (KO) was achieved using CRISPR-Cas9 and the AltR Crispr system (Integrated DNA Technologies). RNA components (IDT) and purified Cas9 (UCB Macrolab) were nucleofected in the SH- SY5Y cell line (Lonza 4D-Nucleofector core unit; see Transfection methods). crRNA sequence and primers used to evaluate cutting are listed in supplementary table 1. Nav1KO clones were identified via sequencing, and the lack of Nav1 in SH-SY5Y cell line was validated via Western Blot.

### Immunostaining

#### Immunocytochemistry

Cell cultures were fixed in 3.7% formaldehyde with 120mM sucrose in phosphate buffered saline (PBS) at 37°C for 15 minutes, then permeabilized with 0.2% Triton-X 100 in PBS for 10min, blocked in 1% bovine-serum albumin in PBS for 30min, and incubated with the following primary antibodies diluted in PBS for 1hr at RT: TuJ1 (1:2000 to stain primary neurons, 1:300 to stain SH-SY5Y cells, Dr. A. Frankfurter, and Neuromics MO15013), anti-Nav1 (generation described above, 1:300), anti-GFP (1:200; Invitrogen A-11122), anti-drebrin (1:400 MBL D029-3), anti-EB1 (1:300, BD Biosciences 610535), anti-cortactin (1:300, Millipore 05-180), anti-tyrosinated tubulin (1:2000; Sigma MAB1864-I), anti-neuron specific enolase (1:300; Novus Biologicals NB110-58870), anti-*α* tubulin (1:1000; abcam ab6161), anti-SMI-31(1:300-1:500; Biolegend 801601), anti-GAP43 1:250; Novus Biological NBP1-92714), anti-nestin (0.5mg/mL; BD Bioscience), anti-GFP (5mg/Ml; Aves Lab GFP-1010); and anti-Tbr1 (1:500, Santa Cruz sc-48816). Immunostaining against Nav1, EB1, fascin or cortactin required 3min fixation with 100% ice-cold MeOH. Following rinsing we incubated with Alexa-Fluor-488, -568, -647- conjugated secondary antibodies, DAPI (1:2000; Biotium 40011), Hoechst (1:1000; Pierce 33258), or phalloidin (1:250; Invitrogen) and in blocking buffer at 37°C for 45min. mVenus or eGFP was used to identify neurons cotransfected with Control or Nav1 shRNA and the mVenus/eGFP signal was enhanced with anti-GFP antibody.

#### Immunohistochemistry

Brain sections were prepared and immunostained according to Castanza *et al* (2021). Briefly, embryonic mouse brain tissues were dissected, fixed in 4% paraformaldehyde (PFA), and were sectioned in a coronal plane at 7 (E12.5) and 12 microns thickness (E14.5) (Figure 1) or 100 mm thickness (E18.5) (Figure 3). No heat induced antigen retrieval (H.I.E.R.) was performed. Sections were blocked for 30 min at room temperature in 10% serum (goat or donkey depending on primary antibody origin species), 3% BSA, and 0.1% Triton X-100 in PBS. Primary antibodies were diluted in blocking solution and incubated overnight at 4°C. Sections were washed 3x5 min in PBS in secondary antibodies were diluted 1:400 in blocking solution and incubated for 2 h at room temperature. Tissue was again washed 3x5 min in PBS, incubated briefly in DAPI when indicated and mounted with Southern Biotech Fluoromount-G (Figure 1) or Vectashield (Figure 3) and allowed to cure before imaging.

### Immunoblotting

Cells were lysed in radioimmunoprecipitation assay (RIPA) buffer (150mM NaCl, 0.1% Triton X-100, 0.5% sodium deoxycholate, 0.1% SDS, 50mM Tris-HCl pH 8.0, with 1mM phenylmethylsulfonyl fluoride in isopropanol added right before lysis) with constant agitation at 4°C for 30 minutes. Lysates were then centrifuged at 12,000 rpm at 4°C for 20min and total protein extracts were boiled in Laemmli sample buffer (BioRad), separated by SDS-PAGE under reducing conditions and transferred to PVDF membranes (Millipore), which were then incubated with the following primary antibodies: anti-GAPDH (1:2000; GeneTex GT239), anti-Nav1 (1:1000), or anti-TrkB (1:500; BD Biosciences 610101). Immunoreactive bands were visualized by the LI-COR Odyssey Imaging System using anti-mouse IRDye 800 and anti-rabbit IRDye 680 (1:5000, LI-COR). To quantify and compare signal intensity, a box was applied to encompass the band of interest, and integrated intensity was determined after background subtraction. In the case of TrkB, we assumed that both bands of the doublet appearing in the SH-SY5Y cell extract corresponded to TrkB, and were included in the boxed region.

### Membrane blebs

Cells were seeded at 263 cells/mm^2^ on 100μg/mL poly-(D)-lysine coated glass coverslips. Rho-kinase Inhibitor Y-27632 (5µM) in DMSO or DMSO alone was applied to Nav1KO SH-SY5Y cells for 2 hours before fixation. SH-SY5Y Nav1KO cells were transfected with 1.5μg of Rac1Q61L or eGFP, and fixed 48 hours after transfection. For quantification of membrane blebs in WT vs Nav1KO cells, cells were plated on coverslips and fixed 24 hours later. A cell was considered to be blebbing if it had at least two blebs in the periphery. For live imaging of membrane blebs, cells were seeded at 263 cells/mm^2^ on 100μg/mL poly-(D)-lysine coated ibidi µ-Plate 24 well plates, and imaged every 30 minutes for 24 hours in an environmental chamber at 37°C and 5% CO_2_.

### Endocytosis labeling

FM4-64 (Thermo T13320) dye application protocol was modified from Bonanomi *et al* 2008. For primary neurons, FM4-64 (10µM) was diluted in Krebs-Ringer-Hepes (KRH) solution, the dye was loaded onto the cells for 1 min, and then washed 2 times with KRH for no more than 2 minutes, and immediately fixed in 3.7% MeOH-free paraformaldehyde (PFA) with 120mM sucrose in PBS for 15min at RT. Acute cholesterol extraction was performed using M*β*CD (5mM, 3 minutes, 37°C), and LY294002 was used to inhibit PI3Kinase (50μm, 30 minutes, 37°C). Images were acquired within 1 week after fixation, as a loss of signal was observed after 1 week. Lysine-fixable dextran (Thermo D1818, 2mg/mL) or transferrin (Invitrogen T23365) was incubated for 10 min at 37°C and immediately fixed according to the MeOH-free PFA protocol. For SH-SY5Y cells, FM4-64 was loaded for 5 minutes, and SH-SY5Y cells were either live imaged or fixed according to the MeOH-free PFA protocol, and fixed cells were incubated with phalloidin (1:100) overnight at 4°C.

### TrkB uptake experiments

TrkB 1D7 (Bai *et al*., 2010), an agonistic monoclonal antibody, was applied in cold HBSS plus 2mM CaCl_2_ to primary neurons for 30 minutes. Samples were washed twice in the cold HBSS solution, and then incubated in 37C HBSS solution for 10 minutes, and immediately fixed with paraformaldehyde for 15 minutes at RT. After fixation, cells were permeabilized with 0.2% Triton-X-100 for 10 minutes, and incubated with Alexa Fluor 568 anti-mouse IgM secondary (1:250, Invitrogen, A-21043)

### Image acquisition

#### Histological tissue sections

For the *in utero* electroporation experiments, images were acquired in 1024 x 1024 mode with a Nikon Ti-E microscope equipped with the A1 laser scanning confocal microscope. We used the following objective lenses (Nikon): 10x PlanApo, NA 0.45 (for images of cortical slices), 60x Apo TIRF, NA 1.49 (analyses the leading process and morphometric analyses of multipolar and bipolar cells). Images were acquired using Nikon software NIS-Elements (Nikon Corporation, Melville, NY).

The immunohistochemistry performed for Nav1 localization was acquired with Zeiss LSM780 confocal microscope, and digital images were adjusted for brightness and contrast in Photoshop (Adobe).

#### Primary neurons

Excluding the endocytosis experiments, fluorescence imaging was carried out using a CSU-X1 spinning disk confocal (Yokogawa) mounted onto an Olympus IX70 microscope and a 20x 0.8 NA Plan APO for shRNA experiments and 60x 1.42 NA Plan APO oil immersion objective for the rest. Fluorescent specimens were excited using a laser launch equipped with the following 50mW solid-state lasers: 405nm, 488nm, 561nm and 640nm. Fluorescence emission was selected through the following band-pass filters: 460/50nm, 525/50nm, 595/50, 700/75. A stack of images was acquired in the z dimension using optical slice thickness of 0.2 using a CoolSNAP HQ2 digital CCD camera (Photometrics) with pixel size of 91 nm.

#### Primary neuron endocytosis and SH-SY5Y experiments

Fluorescence imaging was carried out CSU-X1 spinning disk confocal (Yokogawa) mounted onto a Ti-E microscope with perfect focus system (Nikon) and a 60x 1.4 NA Plan APO oil immersion objective. For the SH-SY5Y neuritogenesis experiment, we used a 20x 0.75 NA objective. Fluorescent specimens were excited using a laser launch equipped with the following 50mW solid-state lasers: 405nm, 488nm, 561nm and 640nm. Fluorescence emission was selected through the following band-pass filters: 460/50nm, 525/50nm, 595/50, 700/75. A stack of images was acquired in the z dimension using optical slice thickness of 0.2 μm using an Photometrics Prime 95B cMOS camera. For the SH-SY5Y neuritogenesis experiment, automatic acquisition was used using the MetaMorph high content screening module.

### Image Analysis

#### Primary neuron neuritogenesis

Morphometric analysis was performed using MetaMorph software (Molecular Devices or Fiji (NIH). Neurons were identified by immunostaining for the neuronal marker ß- III tubulin and neurite length was measured manually using the eGFP signal and computer-assisted analysis from MetaMorph. Neurons without neurites were classified as stage 1, neurons with at least one neurite were considered stage 2 and neurons with one neurite at least twice the length of the other ones was considered the axons and the cells in stage 3.

#### SH-SY5Y neuritogenesis

The length of neurites was measured per field using a custom cell profiler pipeline (available upon request). Briefly, TuJ1 and neuron-specific enolase were used to identify neurogenic cells, expanded DAPI was used to exclude cell bodies in measurements, and a high phalloidin signal was used to exclude non-neurogenic cells that also expressed TuJ1 at low levels. We then calculated total neurite length in the image field divided number of cells (as measured by number of cells identified using DAPI).

#### In utero Nav1 shRNA experiments

The length of the leading process of migrating neurons in the cortical plate and the angle of migration in the Intermediate Zone was measured using Nikon Software NISElements (Nikon Corporation, Melville, NY). The orientation of each leading process was defined as that of a line connecting the cell body center and the base of the leading process and its angle was measured with respect to the pial surface. An average of all angles was used to calculate the deviation of each individual angle to the average. The quantification of bipolar and multipolar neurons was performed by morphological criteria. Briefly, for each neuron in the IZ we quantified the number of non-axon processes (axon was identified by thickness, length and orientation). Neurons were considered bipolar when they presented one axon and one process. Neurons were considered multipolar when they displayed more than one non-axonal process.

#### Nav1 and EB1 analysis

For estimation of the amount of the Nav1 proteins bound to the outer most MT tips; integrated fluorescence intensities within a box of four pixels on a side (outer tip) were measured for each channel after subtracting external background. EB1 centroid was used to position the region of interest.

#### Membrane Ruffles

The F-actin channel was thresholded to encompass the structures. The resulting mask was used to generate a manual selection that would include the F-actin membrane ruffles within the transition zone, identified as the actin-rich region adjacent to the central zone microtubules. From this double selection the area and intensity of the F-actin signal was measured.

#### Endocytosis

Growth cones were identified by their morphology, and a mask was created using the pcs- membrane-ceruleanFP signal for primary neurons, or phalloidin-labeled F-actin for the SH-SY5Y cells. Intensity of endocytosis probes (FM4-64, transferrin, TrkB1D7) was measured in that mask after background subtraction and thresholding. A growth cone was considered to have uptake if there were at least two FM4-64 puncta within the growth cone. For the PI3Kinase treatment, we quantified the percentage of growth cones with FM4-64 uptake, instead of integrated signal intensity, because we unexpectedly observed that LY294002 caused FM4-64 to be taken up all over the cell, not just at growth cones as in controls, thereby complicating intensity measurements.

### Statistical Analysis & Experimental Rigor

All results reported have been observed reproducibly in at least 3 experiments; specific replicate values are indicated in figure legends. Prior to quantitative analysis, sample identity was encoded for blinding of the experimental group. Statistical significance was set at the 95% confidence level (two tailed) and calculated using Prism (Graphpad Software). Values are presented as the mean ± S.E.M.

To assess the normality of the data, we used the D’Agostino-Pearson test to determine deviation or kurtosis. When normality was not met either the Mann-Whitney or Kruskal-Wallis test with Dunn’s multiple comparisons was chosen as described in the figure legends. Otherwise, two-way ANOVA with Bonferroni post-hoc comparisons were used. Pharmacological and genetic experiments were statistically compared to their corresponding control or wild-type counterparts.

## Abbreviations

Nav1: Neuron Navigator 1
BDNF: Brain derived neurotrophic factor
EB(s): end binding protein(s)
MT(s): microtubule(s)
+TIP(s): plus-end tracking protein(s)
F-actin: filamentous actin
GEF: guanine nucleotide exchange factor

## Acknowledgements

We gratefully acknowledge the initial work by Maria-Jose Martinez-Lopez on this project. We thank Halpain lab members Vaidehi Gupta for preparation of neuron cultures, and Eva Desai and Taylor Lichtenberg for their help in image analysis. We are grateful to Franck Polleux for his experimental advice and support of the Nav1 knockdown *in utero* electroporation experiments. We thank Joshua Schwartz and Liz Roberts for advice on CRISPR/Cas9 protocols. We also thank William Mobley and Aaron Johnstone for advice in the design of the TrkB uptake assays, Uri Saragovi and William Mobley for sharing the TrkB 1D7 antibody, Michael W. Davidson for EB3-mcherry, and Carlos Buesa Arjol for supplying various Nav1 constructs. This work was supported by NIH grants NS37311, MH087823, and was a component of the National Cooperative Reprogrammed Cell Research Groups (NCRCRG) to study mental illness and was supported by NIH grant U19MH1 2015-0644.

---

**Supplementary Figure 1.**
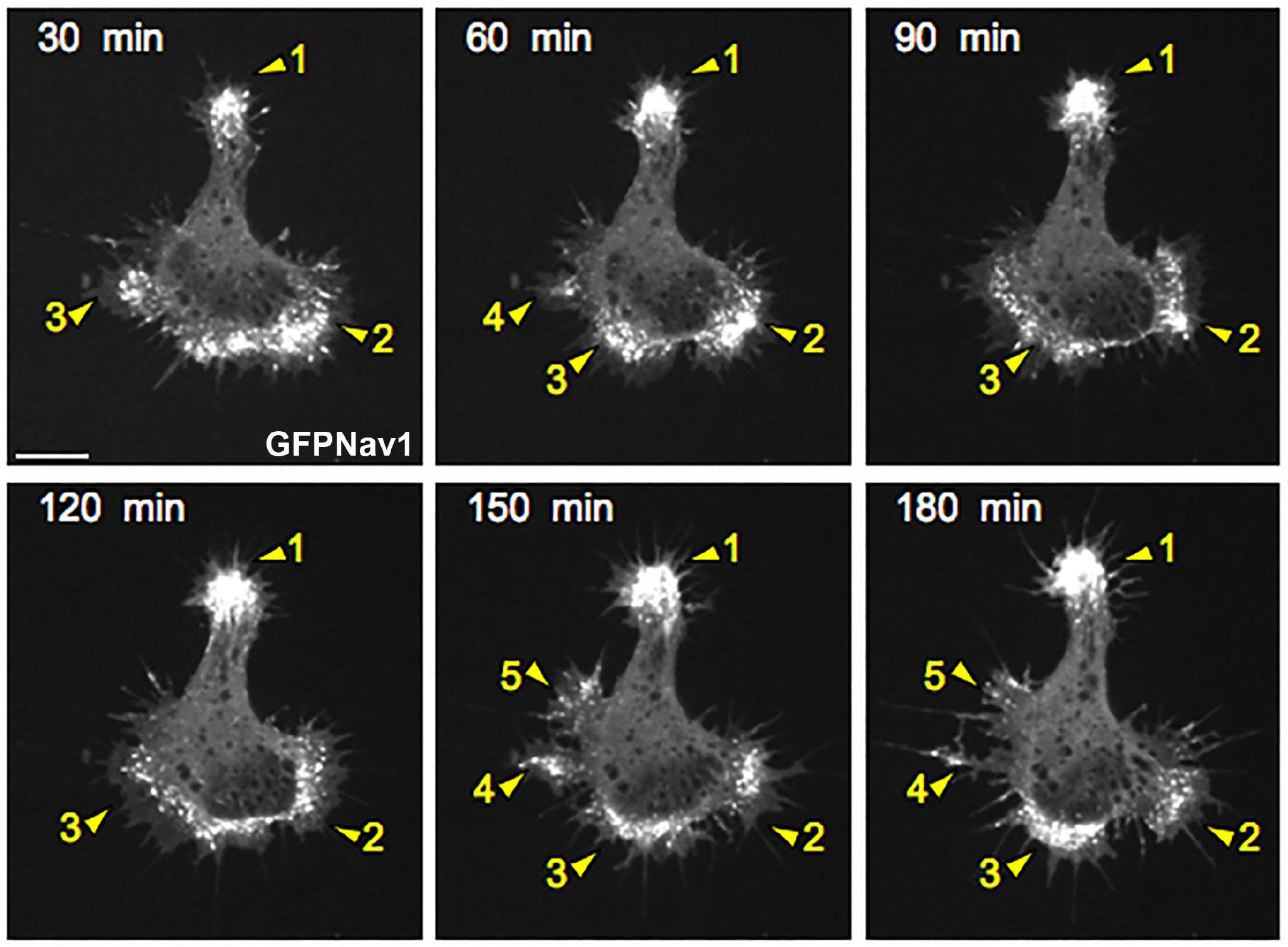
GFP-Nav1 subcellular dynamics during neuritogenesis. Selected frames from a time-lapse sequence showing that Nav1 is enriched in subcellular regions (arrowheads) that extend from the cell body and might commit to becoming neurites. Arrowheads show the appearance of these nascent growth cones over neuronal morphogenesis. Cells presented the characteristic filopodial activity and segmentation of the initial lamellipodia surrounding the soma. Such membrane segmentation commonly precedes the selection of a neurite initiation site in cultured hippocampal neurons. Scale bar = 20μm.

**Supplementary Figure 2.**
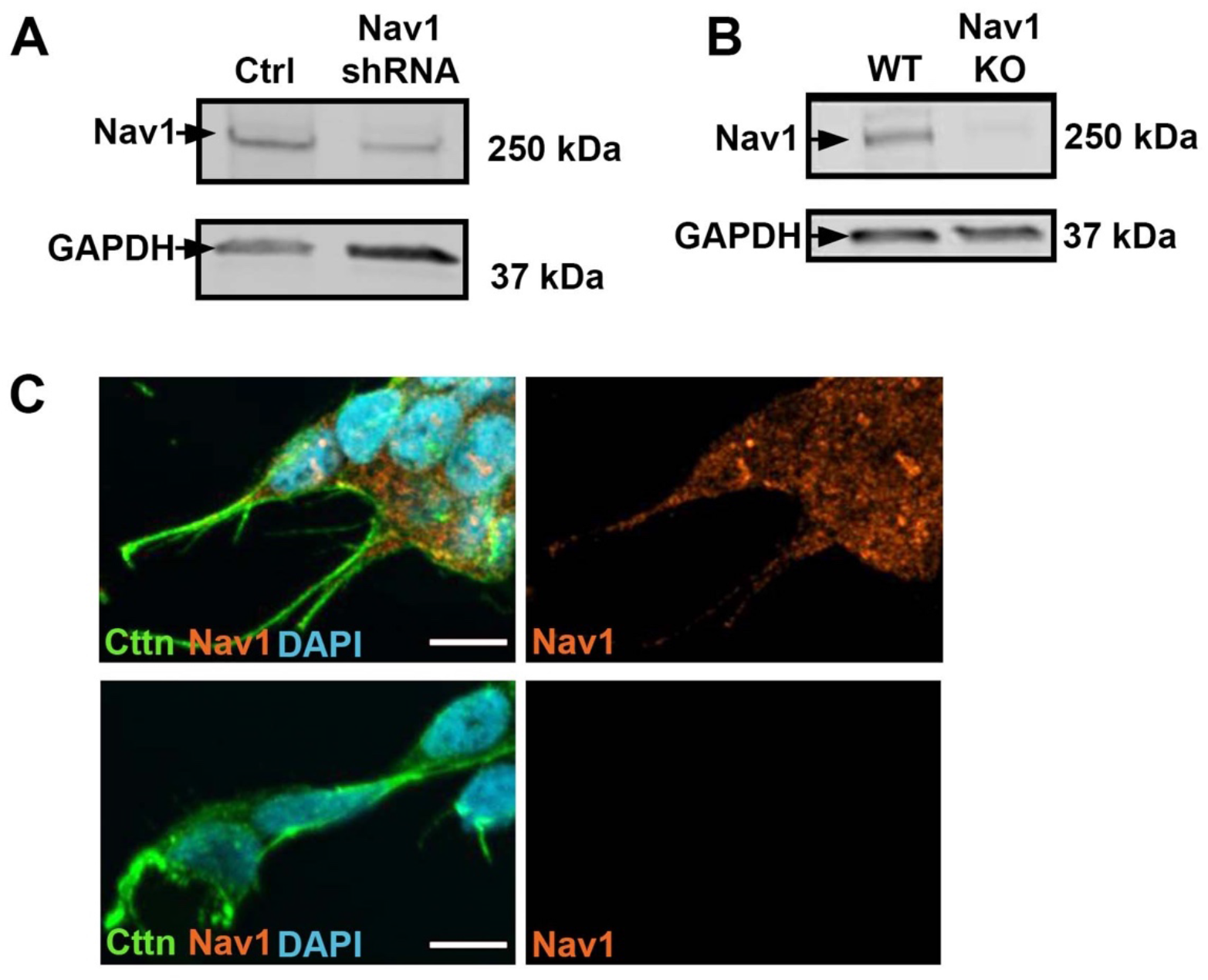
Validation of Nav1 gene silencing and antibody specificity. (A) Immunoblot demonstrating the partial knockdown of Nav1 by shRNA in N1E cells. (B) Immunoblot of WT and Nav1KO SH-SY5Y cells demonstrating the lack of detectable expression of Nav1 expression in cells where the Nav1 gene was silenced via CRISPR/Cas9 gene editing. (C) Representative images of undifferentiated WT and Nav1KO SH-SY5Y demonstrating the lack of detectable Nav1 fluorescence signal in the Nav1KO cells using identical image collection and display settings. Scale bar = 10μm

**Supplementary Figure 3.**
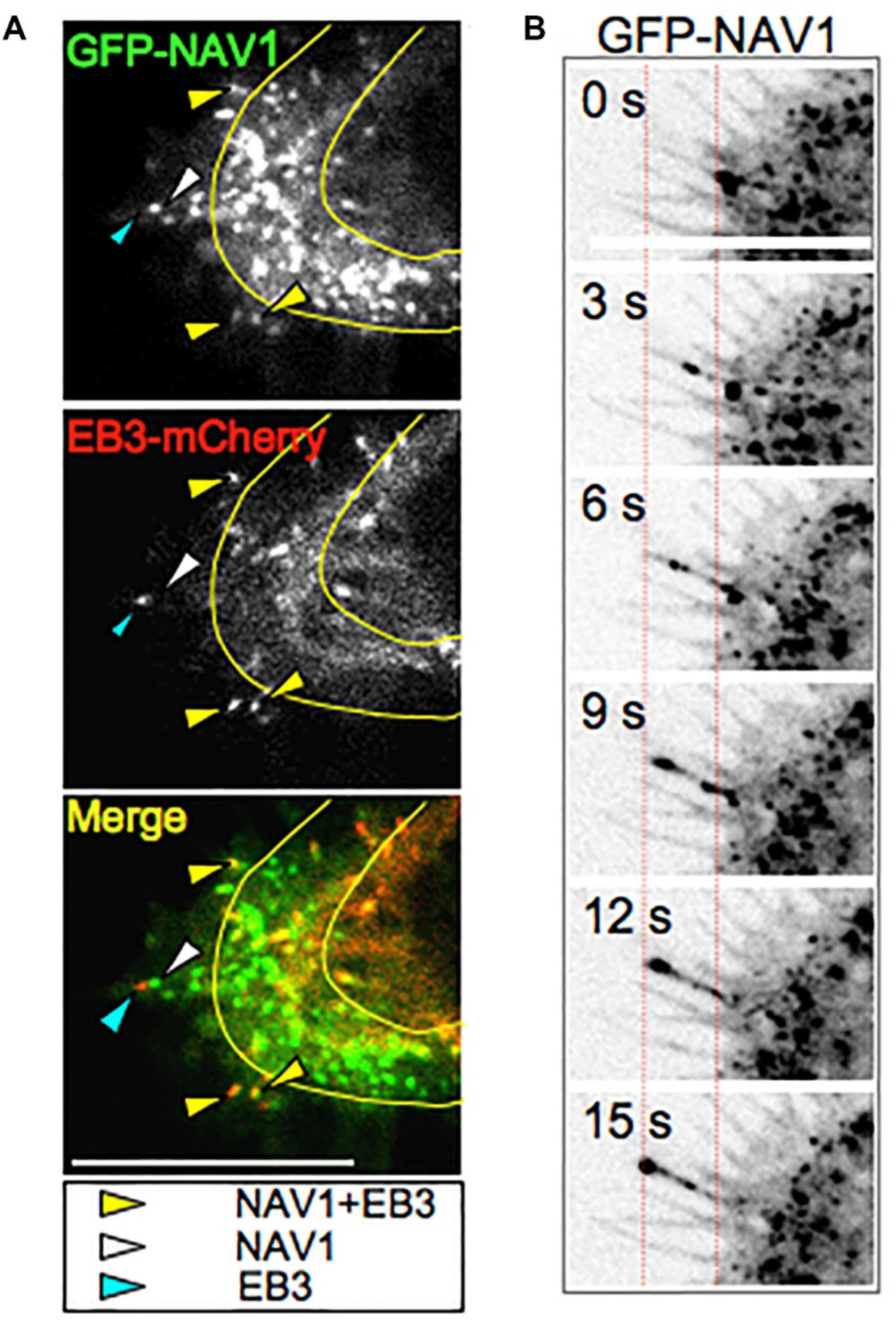
Nav1 associates with some EB1 puncta in the growth cone periphery. (A) Representative frame from a time-lapse sequence of a nascent growth cone cotransfected with GFP-Nav1 and EB3-mCherry. Yellow arrowheads indicate colocalized puncta of GFP-Nav1 and EB3-mCherry; white arrowhead indicates a GFP-Nav1 positive motile punctum lacking detectable EB3-mCherry; cyan arrowhead indicates an EB3-mCherry positive punctum lacking detectable GFP-Nav1. T-zone is highlighted with a yellow line. Scale bar = 10μm (B) Time-lapse sequence of GFP-Nav1 in a nascent growth cone. Red dashed lines indicate the position in the first and last time points of a GFP-Nav1 motile punctum advancing from the T-zone and entering a filopodium in the P-domain. Scale bar = 10μm.

**Supplementary Figure 4.**
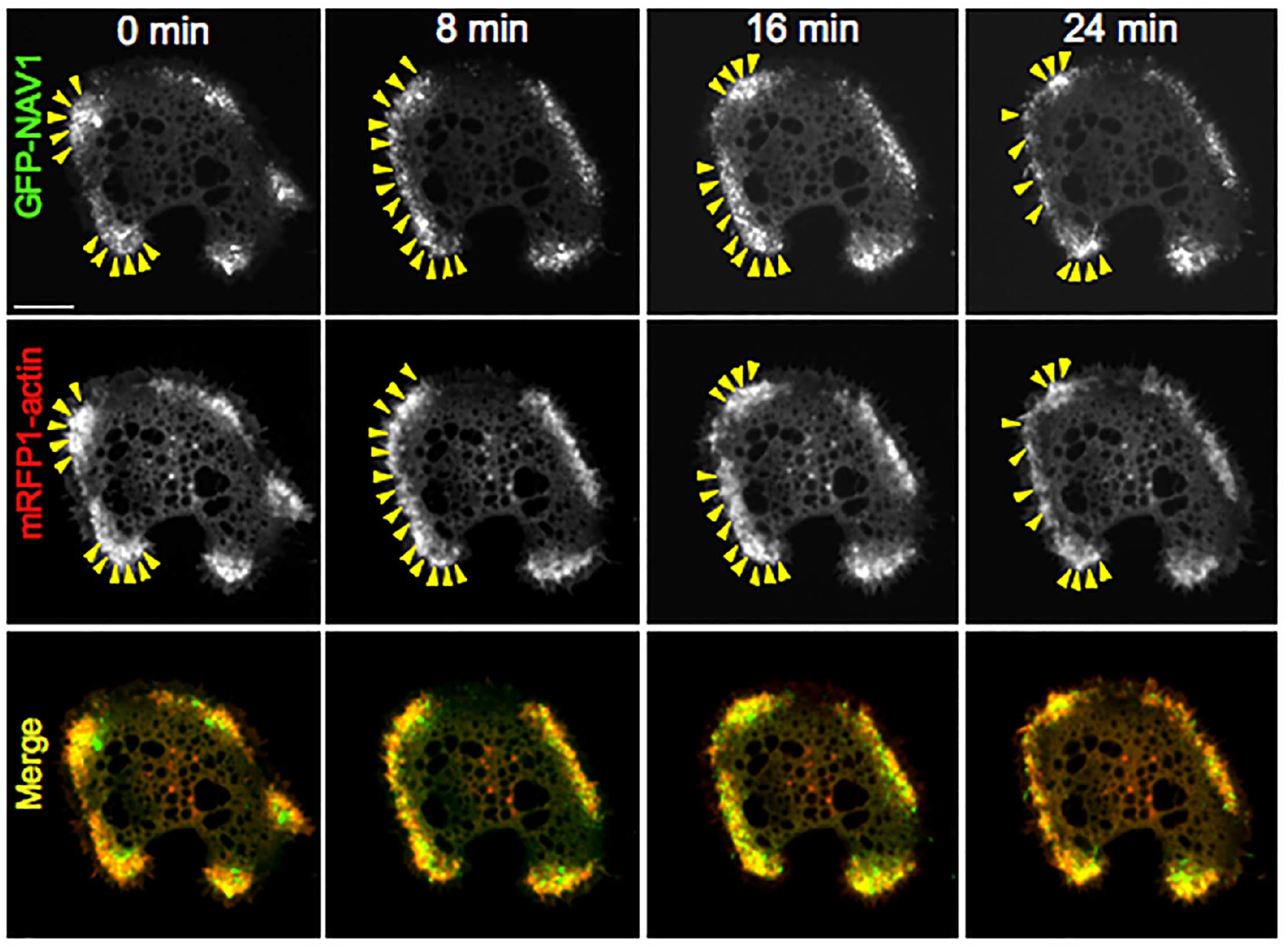
Nav1 and actin colocalize in stage 1 neurons. Time-lapse sequence from a neuron cotransfected with GFP-Nav1and mRFP1-actin; images were acquired every 8 minutes. Note that mobile clusters (arrowheads) containing GFP- Nav1 and mRFP1-actin reorganize in coordinated fashion. Scale bar = 20μm.

**Supplementary Figure 5.**
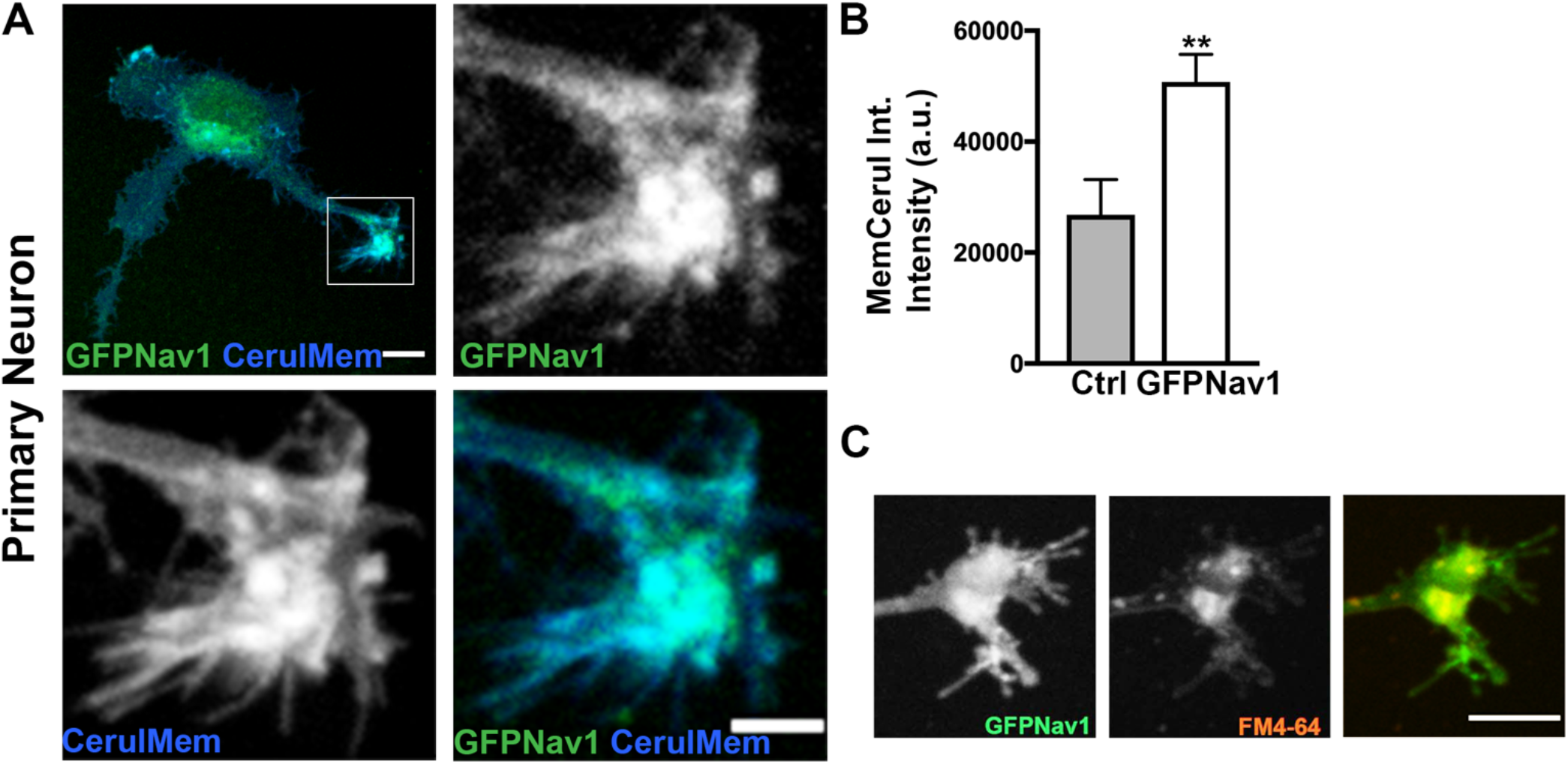
GFP-Nav1 promotes endocytosis and membrane accumulation. (A) Representative image showing growth cone showing enriched pcs-membrane-cerulean-FP in the same growth cone areas where GFP-Nav1 is enriched. Scale bars = 10μm for large image, 5μm for zoomed image. (B) GFP-Nav1expressing growth cones have significantly higher intensity of cerulean-tagged membrane than control growth cones. Statistics: Mann-Whitney, ***p<0.0001, n = 4 experiments; Control = 52 growth cones, GFP-Nav1 = 142 growth cones GFP-Nav1. (C) Representative image showing FM4-64 is taken up in GFP- Nav1-expressing WT SH-SY5Y growth cone. Scale bar = 10μm.

**Supplementary Table 1.**
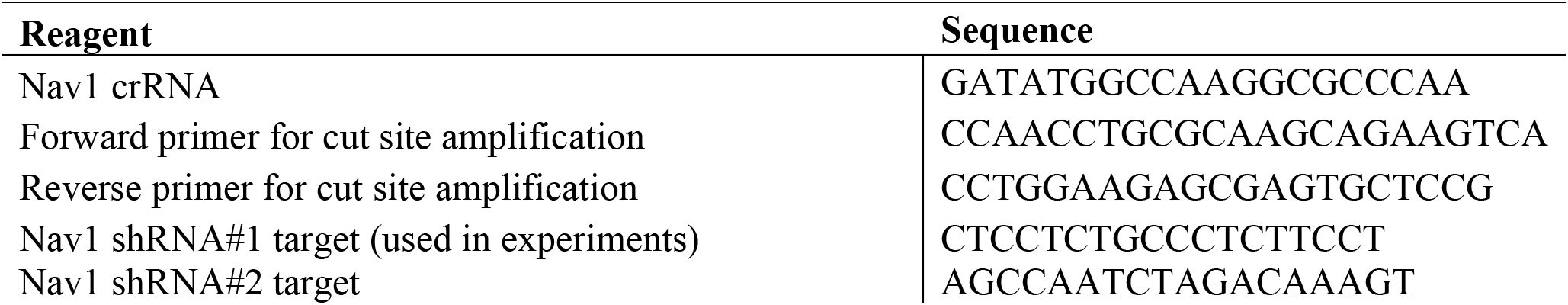
List of target or primer sequences.

